# Cultivation of SAR202 Bacteria from the Ocean

**DOI:** 10.1101/2023.03.25.534242

**Authors:** Yeonjung Lim, Ji-Hui Seo, Stephen J. Giovannoni, Ilnam Kang, Jang-Cheon Cho

## Abstract

Here we report the first successful cultivation of SAR202 bacteria, a superorder in the phylum *Chloroflexota*, which have long been at the top of “most wanted” lists of uncultivated microbial life. It has been proposed that ancient expansions of catabolic enzyme paralogs in SAR202 broadened the spectrum of organic compounds they could oxidize, leading to transformations of the Earth’s carbon cycle. We cultured the cells from surface seawater using dilution-to-extinction culturing. Their growth was very slow (0.18-0.24 day^-1^) and was inhibited by exposure to light. The genomes, of ca. 3.08 Mbp, encoded archaella, archaeal motility structures, and multiple sets of paralogs, including 80 genes in enolase superfamily and 44 genes in NAD(P)-dependent dehydrogenase family. We propose that these paralogs participate in multiple parallel pathways of non-phosphorylative sugar and sugar acid catabolism, and demonstrate that, as predicted by this scheme, the sugars ʟ-fucose and ʟ-rhamnose and their lactone and acid forms are utilized by these cells.

## Main

The SAR202 clade in the phylum *Chloroflexota* is ubiquitously distributed in the ocean, accounting for 10–30% of planktonic prokaryotes in the deep sea^1–7^. Various properties associated with organoheterotrophy and sulfur and nitrogen metabolism have been interpreted from SAR202 metagenome assemblies and single cell genome sequences^6, 8–12^. The seven groups (subclades) of SAR202 are individually distinct in the numbers and types of paralogs they contain^8, 12^, suggesting a relationship between the paralog evolution and niche specialization of subclades.

Paralogous flavin-dependent monooxygenase genes in group III SAR202, in some cases exceeding 100 per genome, are proposed to have evolved to harvest carbon and energy from diverse organic molecules that accumulated in the oceans during the expansion of the Earth’s carbon cycle, following the rise of oxygenic phototrophy^8, 12, 13^. Similarly, in group I SAR202, large expansions of paralogs in the enolase protein superfamily are proposed to have evolved to enable these cells to metabolize compounds that resist biological oxidation because of their chiral complexity. The early branching of these paralogs in phylogenetic trees indicates that SAR202 cell evolution was a crucible for their diversification^8, 12, 13^.

SAR202 cells are found throughout ocean water columns, reaching highest numbers near the ocean surface, but they contribute a higher percentage of all plankton cells in the meso-, bathy-, hadal-, and abyssopelagic^5, 10, 12^. Groups I and II have rhodopsin genes in their genomes and are the most abundant ones in epipelagic environments, whereas group III is largely responsible for the high relative abundance of SAR202 in the dark ocean.

The cultivation of unrepresented cell types is a priority for microbiologists because cells frequently exhibit properties that cannot be easily predicted from their genomes^14^. A recent study has shown that genome-based inference fails to reliably predict catabolic pathways for more than 50% of carbon sources utilized by diverse prokaryotes that have well-curated phenotypic data^15^. Despite keen interest and technological advances, SAR202 and many other important taxa remain on “most wanted” lists of uncultivated cell types^14, 16, 17^.

Here we report the first successful cultivation of SAR202 bacteria. Twenty-four isolates of subclade I were retrieved from surface seawater samples by dilution-to-extinction in sterile seawater media. Metabolic reconstruction supported by experimental data with cells implicated enolase and dehydrogenase paralogs in non-phosphorylative sugar oxidation. We propose that multiple parallel metabolic pathways of this type enable these cells to harvest complex mixtures of sugar-related compounds from dissolved organic carbon pools. SAR202, which are found throughout the water column of modern oceans, evolved concurrently with the rise of oxygenic phototrophy^13^. We propose they expanded into the niche of harvesting dilute and diverse carbohydrate-related molecules as the oceans are oxidized.

## Results and Discussion

### The first successful cultivation of the SAR202 clade

Dilution-to-extinction experiments with low-nutrient heterotrophic media (LNHM; Extended Data Table 1) in microtiter dishes retrieved twenty-four SAR202 group I isolates from surface samples (depth, 10 m) from two stations (GR1 and GR3) located nearby Garorim Bay of the Yellow Sea (Extended Data Fig. 1a). Four conditions, differing by catalase addition and light exposure (continuous dark vs. 14:10 h light-dark cycle) yielded 610 strains, of which 24 belonged to the SAR202 clade (Extended Data Table 2). All 24 SAR202 strains were obtained from cultures incubated continuously in dark (Extended Data Table 2).

Phylogenetic comparisons showed that the isolates had nearly identical 16S rRNA gene sequences, were affiliated with the SAR202 group I (Fig. 1a)^3, 8, 12^, and corresponded to major (greater than 68%) amplicon sequence variant (ASV) of the SAR202 clade (0.7–1.0% in total prokaryotes) in the water samples (Extended Data Fig. 2).

**Fig. 1.**
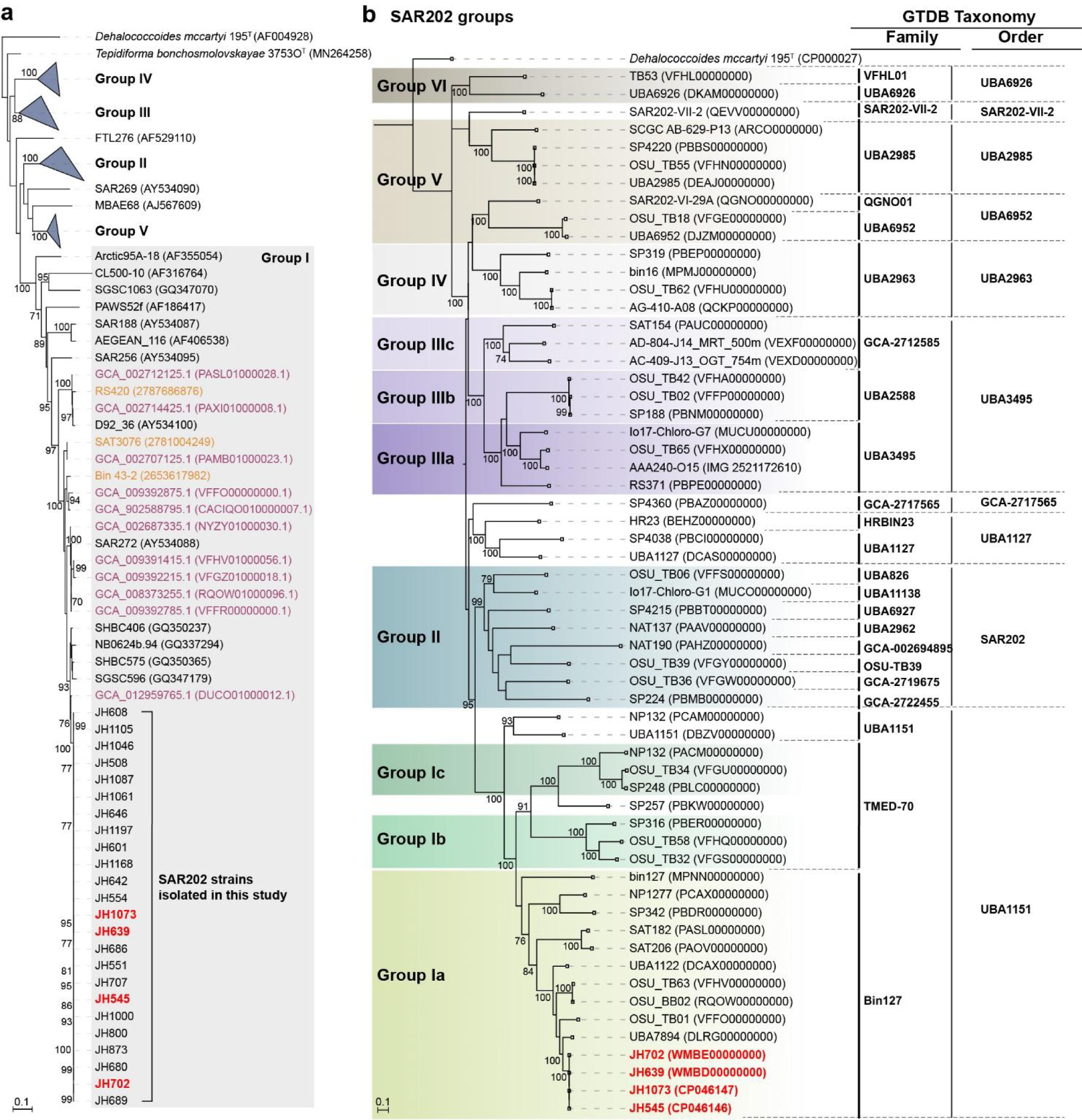
Phylogenetic position of SAR202 strains isolated in this study. **a,** Phylogenetic tree based on 16S rRNA gene sequences showing the relationship among the 24 SAR202 strains isolated in this study and closely related sequences retrieved from various databases, including GTDB (marked in purple), IMG (yellow), GenBank, and EzBioCloud. The four SAR202 strains selected for genome sequencing are marked in red. RAxML (v8.2.12) was used for tree building with GTRGAMMA model. Bootstrap supporting values (≥70%; from 100 resamplings) are indicated. *Dehalococcoides mccarty* was set as an outgroup. Bar, 0.1 substitutions per nucleotide position. **b,** Phylogenomic tree of the SAR202 clade. The four genomes of SAR202 strains from this study (marked in red) and other MAGs and SAGs from previous studies were classified using GTDB-Tk. Species cluster-representative genomes of the GTDB taxa, into which the analyzed SAR202 genomes were classified, were included for tree building. Designation of the SAR202 subgroups following the classification scheme of a previous study^12^ is indicated on the left side of the tree. On the right side of the tree, taxonomic assignment of the genomes at the family and order levels according to the GTDB (release 202) is shown. The tree building was performed using RAxML (v8.2.12), with PROTGAMMAAUTO option, based on a concatenated alignment of core genes obtained by UBCG pipeline. Bootstrap supporting values (100 iterations) are indicated on the nodes. Bar, 0.1 substitution per amino acid position.

### The growth of SAR202 isolates was very slow and inhibited by light

Four strains selected for further experiments, JH545, JH702, JH639, and JH1073, behaved similarly, growing slowly (0.18–0.24 day^-1^) and displaying sensitivity to light (Fig. 2a–c). Strain JH545 grew optimally at 15–20 ℃ (Extended Data Fig. 3), and therefore all subsequent experiments were performed at 20 ℃. Approximately 50 days were required to reach stationary phase at maximum cell densities of ∼2 × 10^8^ cells mL^-1^ (Fig. 2a).

**Fig. 2.**
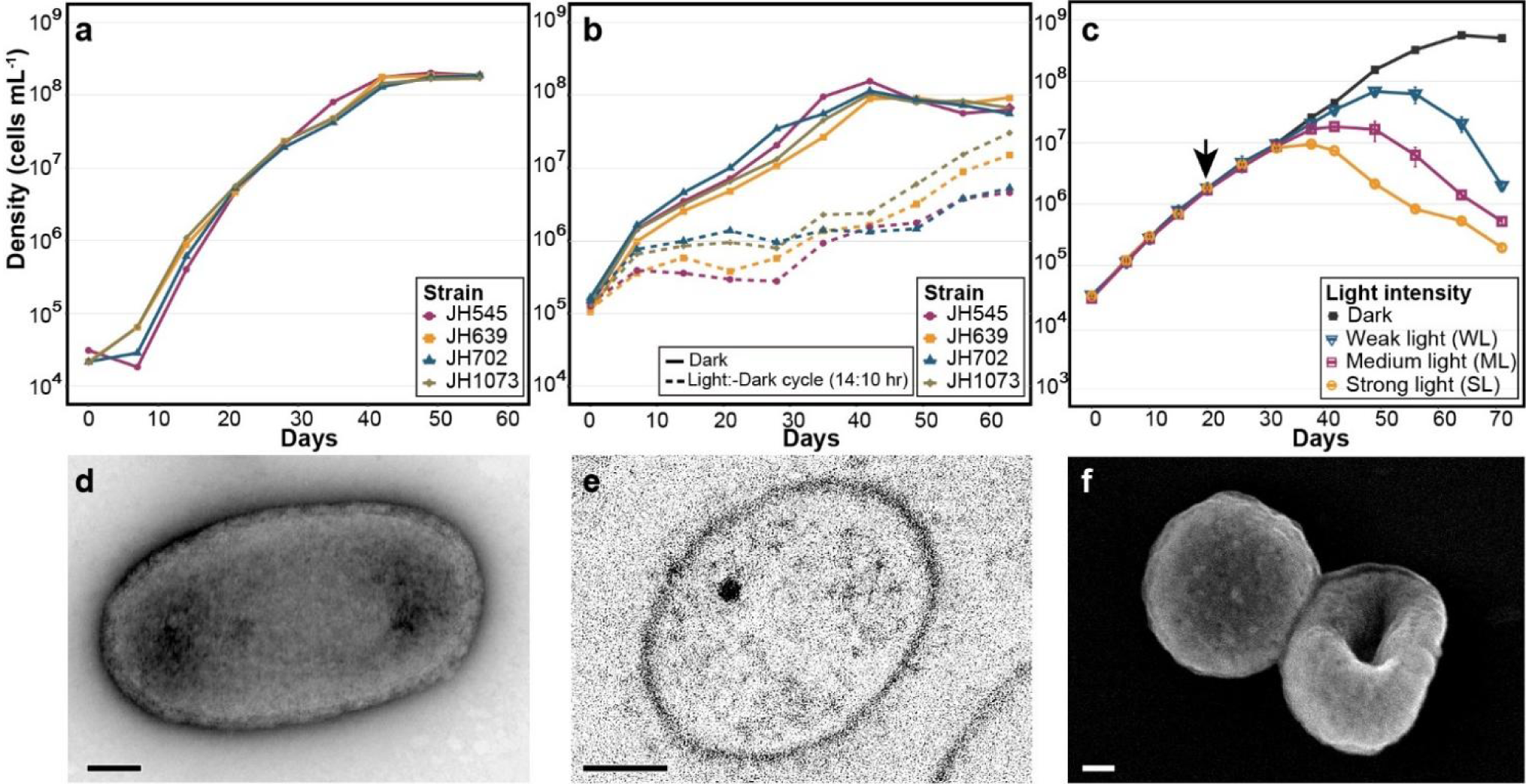
Growth and morphology of SAR202 isolates. **a,** Initial growth curves of four representative SAR202 strains in low nutrient heterotrophic medium (LNHM) under dark condition. **b,** Growth curves of the four strains under continuous dark and light-dark conditions in LNHM. Light-dark cycle (14:10 h) was applied using LED lamps with a light intensity of ∼155 μmol photons m^⁻2^ s^⁻1^. For the dark condition, tubes were wrapped in aluminum foil. **c,** Growth curves of strain JH545 under dark condition and various light intensities in LNHM5x. All cultures were initially incubated in dark (*i.e.*, the tubes were wrapped in aluminum foil). At the day indicated by a black arrow, some cultures were shifted to light condition (*i.e.*, the aluminum foil was removed) and exposed to light with the following intensities: WL, ∼45; ML, ∼89; SL, ∼134 μmol photons m^⁻2^ s^⁻1^. Experiments were performed in triplicates. Error bars indicate standard deviation. Note that the error bars are hidden when they are shorter than the size of the symbols. **d–f,** Electron micrographs of strain JH545 cells observed by TEM (**d**), thin-section TEM (**e**), and SEM (**f**). Scale bars, 100 nm.

The growth of all four strains was inhibited by light-dark cycles with broad spectrum LED lights (Fig. 2b), and experiments with strain JH545 showed that continuous exposure to light caused growth inhibition followed by cell death at all light intensities tested (∼45–134 μmol photons m^-2^ s^-1^), with variations in response levels (Fig. 2c).

Although the isolates were pure cultures, TEM and SEM microscopy showed short rods (∼0.8 × 0.4 μm), cocci (diameter, ∼0.5 μm), discs, and discs with biconcave centers that sometimes appeared to be toroidal, as reported for the strains of *Dehalococcoides*^18, 19^, a genus belonging to the same class (*Dehalococcoidia*) as SAR202 (Fig. 2d–f and Supplementary Fig. 1). Thin-section TEM images indicated monoderm cell envelopes, similar to other *Chloroflexota* (Fig. 2e)^20, 21^.

### Genomic features

#### General genome features and phylogenomics

The genomes of the four SAR202 strains were 3,083–3,094 kb in length with 51.8% GC content, ∼87.5% coding density, and at least 99.9% average nucleotide identity (ANI) among the strains. The two genomes sequenced on PacBio platform (JH545 and JH1073) were assembled into one circularly closed contig, whereas the other two genomes sequenced with Illumina technology were composed of more than 30 contigs (∼846 kb of N50 for both genomes) (Extended Data Fig. 4c). The circular map of the JH545 genome showed several regions with anomalous GC content and GC skew (Extended Data Fig. 4a), many of which overlapped with the predicted genomic islands (Extended Data Fig. 4b), suggesting the influence of horizontal gene transfer.

The four genomes shared similar features, leading us to focus on one of them, JH545, for further analysis (Fig. 3). In accord with previous studies, the genome annotation indicated central carbon and energy metabolism typical of aerobic organoheterotrophs, with some genes indicating capacities for lithotrophy by sulfide oxidation (sulfide:quinone oxidoreductase) and anaerobic respiration by nitrate reduction (NapAB) and N_2_O reduction (NosZ) (Supplementary Text). The presence of NapAB and NosZ has been reported in the SAR202 MAGs obtained from the northern Gulf of Mexico “dead zone”, where the expression of these genes was detected in the samples with lowest dissolved oxygen^9^. In agreement with previous genome analyses of *Dehalococcoidia* members^19, 22, 23^, peptidoglycan biosynthesis was not encoded in the SAR202 genomes. The TEM images showed a layer outside of the cell membrane of JH545 (Fig. 2d–e), reminiscent of the S-layer observed in peptidoglycan-lacking *Dehalococcoidia* strains^24, 25^.

**Fig. 3.**
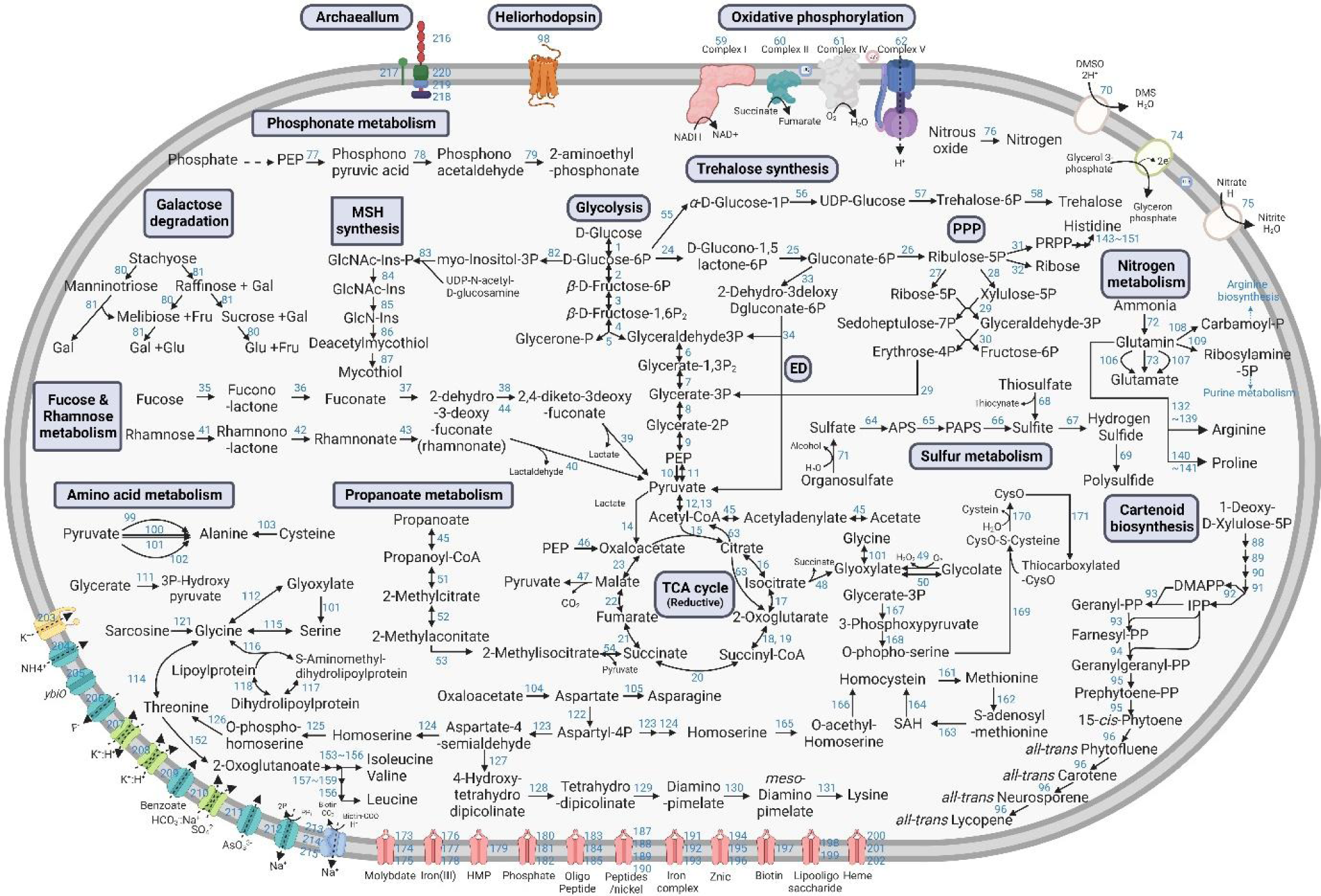
Metabolic pathways of strain JH545. Some key metabolic pathways are indicated by blue boxes. Detailed information on the enzymes corresponding to the numbers next to arrows is available in Supplementary Table 3. The figure was created with BioRender (https://biorender.com/).

Whole genome phylogenies of the Genome Taxonomy Database (GTDB) indicate that the SAR202 group I cells we cultured are the first isolates of an uncultured order (UBA1151) within a monophyletic superorder comprised of all SAR202 (Fig. 1b and Supplementary Text). We propose the provisional taxonomic name “*Candidatus* Lucifugimonas marina”, which includes strains JH639, JH702, and JH1073, in addition to strain JH545 as the type strain. To accommodate this novel genus and species, we also propose the family “*Candidatus* Lucifugimonadaceae” fam. nov. and the new order “*Candidatus* Lucifugimonadales” ord. nov. within the class *Dehalococcoidia* (Supplementary Text).

#### Archaellum

A gene cluster for archaellum, an archaeal motility structure, was predicted in the four SAR202 genomes. This gene cluster is similar to conserved arrangements of archaellum genes observed in archaea^26, 27^ and includes six tandem copies of *flaB* (encoding archaellin), followed by the genes *flaGFHIJ* (Extended Data Fig. 5). Seven additional copies of *flaB* genes were scattered throughout the genome. A candidate gene for FlaK, a family of prepilin peptidases that remove signal peptides from archaellin^26^, was also predicted based on the assignment to COG1989 and the presence of domain PF01478 (Peptidase_A24; Type IV leader peptidase family)^28^.

Archaella are related to type IV pili and have no evolutionary relationship to bacterial flagella. This is the first report of an archaellum gene cluster in bacterial isolates, although a *Chloroflexota* MAG from aquifer sediment was previously reported to harbor a gene cluster for archaellum^29^. Searches for FlaB homologs in the IMG database revealed that two SAR202 MAGs from the Gulf of Mexico had archaellum gene clusters very similar to that of JH545 (Extended Data Fig. 5)^9^. Exploration of FlaB (K07325) distribution using the AnnoTree database showed that most (45 of 52) bacterial genomes harboring *flaB* were affiliated to *Chloroflexota*. The remaining seven bacterial genomes having *flaB* were distributed among seven phyla, indicating that this gene is very rare among other bacteria. Of the 45 *flaB*-harboring *Chloroflexota* genomes, 40 were among the 212 *Dehalococcoidia* genomes in the database, and the remaining 5 were in *Anaerolineae*. This distribution suggests that archaella genes might have been transferred from *Archaea* to an ancestor of *Chloroflexota* and retained in the class *Dehalococcoidia*. Investigation on the 45 *Chloroflexota* genomes having FlaB (found in AnnoTree) and BlastP searches at NCBI (FlaB of strain JH545 as a query) revealed that at least two *Chloroflexota* strains harboring archaella gene cluster (flaB and several other genes) have been cultivated: *Litorilinea aerophila*^30, 31^ and *Aggregatilinea lenta*^32^ (Extended Data Fig. 5). No archaella were observed during microscopic examination of the SAR202 cells.

#### Heliorhodopsin

Two copies of heliorhodopsin (HeR) genes were detected in the JH545 genome. In a recent study of marine SAR202 MAG/SAGs, rhodopsin genes were found in 28 group I and II genomes, all retrieved from water depths of less than 150 m. An HeR gene was reported in a single group II genome^12^. Therefore, this is the first report of HeR in SAR202 group I. The two HeR copies found in the JH545 genome exhibited ∼83% amino acid identity and were located one gene downstream of DNA photolyase, which repairs DNA damage caused by UV exposure using visible light (Extended Data Fig. 6a). Multiple sequence alignment showed that the two copies differed in the amino acid residue that caused a spectral shift in an Ala scanning mutagenesis study^33^ (W163 in 48C12; Extended data Fig. 6b), suggesting that the two HeR copies might have different absorption spectra. Although the function of HeR remains unresolved, it has recently been suggested that it might function as a light sensor that regulates responses to light-induced oxidative stress^34, 35^.

#### Transporters and sulfatases

In accord with a previous study^9^, a large number of major facilitator superfamily (MFS) transporters were found in the JH545 genome, including 42 proteins assigned to COG0477 (Extended Data Table 3). MFS is a very large family of membrane transporters that are known to transport a variety of compounds, including mono-and oligo-saccharides, amino acids, and nucleosides^36, 37^. It is noteworthy that some substrates that we report below enhance the growth of JH545 (see the next section on COG4948) are known to be transported by MFS proteins^38–40^.

Eighteen proteins in the JH545 genome were assigned to COG3119, annotated as arylsulfatase A or a related enzyme (Extended Data Table 3). Sulfatase paralogs have been reported previously in SAR202 groups I and II^12^. Arylsulfatases catalyze the desulfation of sulfated carbohydrates in some catabolic pathways, for example the degradation pathway of ulvan by a marine bacteria^41^.

### Genomes of SAR202 group I have the highest proportion of COG4948 paralogs among all prokaryotes, and these paralogs are highly divergent

The genomes of cultivated SAR202 we report encoded 80 COG4948 proteins in the mandelate racemase family within the enolase superfamily (∼2.8% of CDS; Extended Data Table 3). Many marine bacteria have minimal genomes containing few paralogs, and thus the very large sets of paralogs present in SAR202 genomes attracted attention when they were first noticed^12^.

We sought to establish whether the expansion of COG4948 in SAR202 group I is unusual in the context of prokaryotic diversity. We analyzed the genomes in GTDB, one of the most phylogenetically comprehensive genome databases. Because COG annotation is not scalable, we counted the numbers of the two Pfam domains (PF02746 and PF13378) corresponding to COG4948 (see Materials and Methods for details) and calculated the proportion of COG4948 proteins among all CDSs in each genome. Among the 47,894 species cluster-representative genomes of GTDB (R202), all 48 SAR202 group I (o UBA1151 in GTDB) genomes ranked in the top 66 except one that ranked the 134^th^, in proportions of COG4948 (Fig. 4a), demonstrating that the paralog expansion of COG4948 is a prominent feature of SAR202 group I across all prokaryotes.

**Fig. 4.**
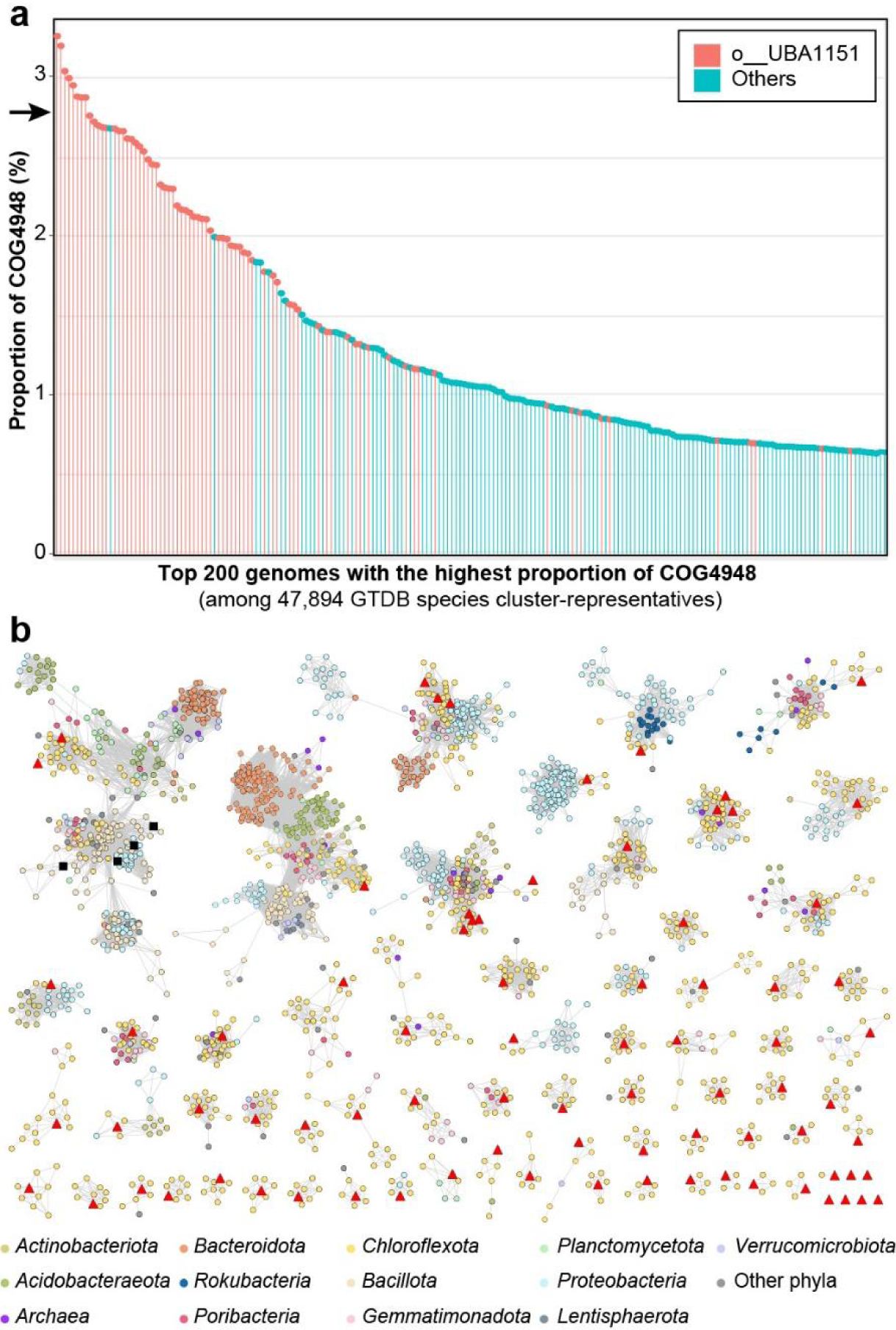
High proportion and diversity of COG4948 proteins in the SAR202 clade. **a,** Proportion of COG4948 proteins among the species cluster-representative genomes of GTDB (release 202; 47,894 genomes). Only the top 200 genomes with the highest proportion are included. Genomes of o_UBA1151 (SAR202 group I) are indicated with a distinct color. All 48 o UBA1151 genomes are included in this figure owing to the high proportion of COG4948. The black arrow on the y-axis indicates the proportion of the four SAR202 genomes sequenced in this study (∼2.8%). **b,** SSN (sequence similarity network) analysis of diverse COG4948 proteins predicted in the JH545 genome. UniProt proteins with the two Pfam domains, PF02746 and PF13378, were included in the analysis together with the 80 COG4948 proteins of strain JH545. Only the protein clusters including the JH454 proteins were retained in this figure after applying a threshold cutoff value to the alignment score (≥150). Nodes (proteins) are colored by their phylum-level taxonomy, following the color codes at the bottom. The 80 COG4948 proteins of the JH545 genome are indicated in red triangles. The four proteins that have been studied experimentally are indicated with black rectangles (UniProt ID: D8ADB5, C9A1P5, C6CBG9, and A8RQK7).

We analyzed sequence divergence among the 80 COG4948 proteins of the JH545 genome by building a sequence similarity network (SSN) using the Enzyme Function Initiative’s Enzyme Similarity Tool (EFI-EST)^42^. Most of the ∼70 clusters that contained JH545 proteins as their members included only one JH545 gene, indicating that the JH545 COG4948 proteins are highly divergent (Fig. 4b). While many of the largest COG4948 clusters included diverse bacterial phyla, many small clusters were comprised mainly of *Chloroflexota*, suggesting that the COG4948 protein family, which is distributed widely across prokaryotes, diversified in *Chloroflexota*. Only the largest cluster, which included two JH545 proteins, contained biochemically studied proteins^43^ (Fig. 4b).

### Abundant COG paralogs of SAR202 participate in the degradation of sugars, lactones, and sugar acids

We propose that seven sets of paralogs (COG1028, COG0667, COG3618, COG4948, COG1063, COG3836, and COG0329), including the most abundant COG4948 paralogs, act concertedly in parallel pathways that harvest energy from diverse carbohydrates (Fig. 5b), and we provide experimental evidence that cultured SAR202 group I cells respond to predicted substrates of these pathways (Fig. 5a). The largest COG sets in the SAR202 genomes, in addition to the mandelate racemases (COG4948), included NAD(P)-dependent dehydrogenases (COG1028), aldolases (COG3836), and oxidoreductases (COG0667; Extended Data Table 3). These enzymes are found in non-phosphorylative pathways of various sugars and their acid catabolism (*e.g.*, fucose, rhamnose, arabinose, and xylose)^39, 44–49^. Central to these pathways is the pairing enzymes from the two largest paralog sets, mandelate racemase-like enzymes and NAD(P)-dependent dehydrogenases, which catalyze a dehydration reaction followed by an oxidation reaction resulting in a flow of electrons for respiration. A previous SAR202 pangenome analysis of variation in COG copy number found significant positive correlations in the numbers of copies of the same seven COGs (Extended Data Table 3) in SAR202 group I genomes relative to other SAR202 genomes^12^. The phylogenetically correlated distributions of these paralog expansions support our prediction that they play coordinated metabolic roles.

**Fig. 5.**
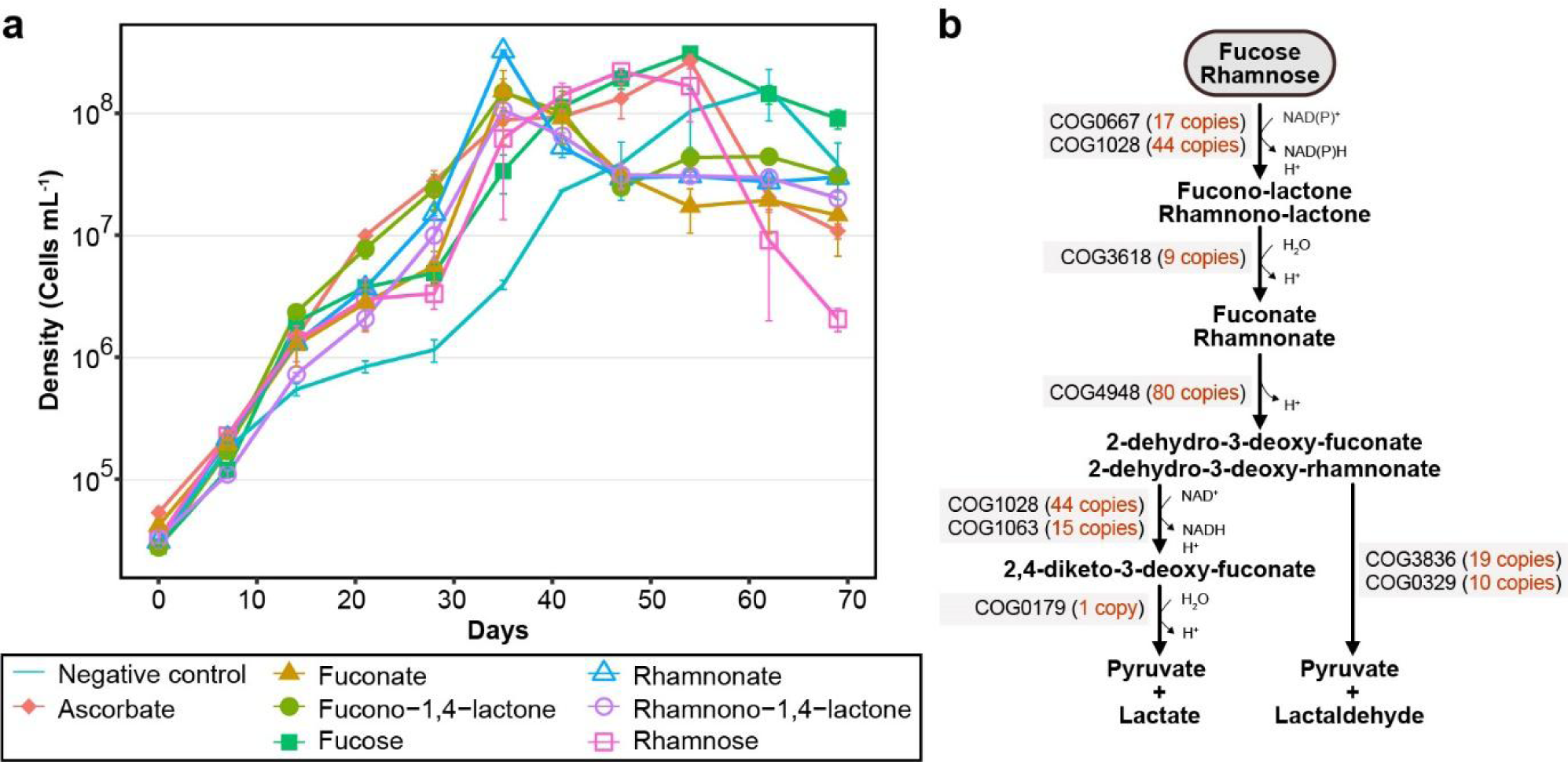
Growth curves of strain JH545 with various carbon compounds and putative non-phosphorylative catabolic pathways of L-fucose and L-rhamnose. **a,** Growth of strain JH545 on various carbon compounds that were predicted to require COG4948 enzymes for metabolization. Carbon compounds were added to the ASW5x media (Extended Data Table 1) at a final concentration of 250 μM. The incubation was performed in dark at 20 ℃. All experiments were performed in triplicates. Error bars indicate standard deviation. Note that the error bars are hidden when they are shorter than the size of the symbols. Negative control, no additional carbon compounds were added. **b,** Putative non-phosphorylative catabolic pathways of L-fucose and L-rhamnose inferred from the SAR202 genome data. The number of proteins assigned to the COGs are indicated within parentheses next to the COG IDs at each step.

We reconstructed non-phosphorylative pathways for the oxidation of two sugars relevant to marine environments, fucose and rhamnose^50, 51^, in the genomes of the cultured SAR202 strains based on several previous studies^46, 47, 52–55^ (Fig. 5b). Kinase-dependent phosphorylative pathways for fucose and rhamnose catabolism were not found in the genomes. In the pathway reconstruction, L-fucose and L-rhamnose share the same COG annotation at each step, yielding pyruvate and lactate or lactaldehyde as final products. Seven of the eight COGs represented in the reconstructed pathways for fucose and rhamnose catabolism were from the abundant paralog sets described above, ranging from 9 to 80 variants of each COG (Extended Data Table 3).

Because pyruvate and reduced nucleotide cofactors are produced by these pathways, L-fucose and L-rhamnose were predicted to serve as carbon and energy sources. We tested the growth response of strain JH545 to ʟ-fucose, ʟ-rhamnose, and their lactone and acid forms. Ascorbate was also tested because an ascorbate degradation pathway requiring a COG4948 enzyme^39, 48^ was nearly complete in the JH545 genome (Extended Data Fig. 7). The growth of strain JH545 in artificial seawater media was enhanced by all substrates tested (Fig. 5a). Growth rates and cell densities in late exponential phase (∼35 days of incubation) were more than 10 times higher in substrate-amended cultures, although the cultures reached similar densities in stationary phase.

### SAR202 isolates represent a cell type that is common in the euphotic zone but relatively more abundant in the dark ocean

The vertical distribution of JH545 in marine metagenomes followed patterns previously observed for several SAR202 group I members^12^. In metagenomes from *Tara* Oceans and station ALOHA, the relative abundance of JH545 DNA, as measured by RPKM (the number of recruited reads per kilobase of genome per million total reads), was higher in the mesopelagic zone (200–1,000 m) compared with the euphotic zone (surface to 200 m) (*P* < 10^-6^; Mann–Whitney *U* test; Fig. 6c). In metagenomes from several marine trenches, the relative abundance of JH545 also increased with increasing depth from euphotic zone to abyssopelagic zone (4,000–6,000 m) through mesopelagic and bathypelagic (1,000–4,000 m) zones (Fig. 6a). But, in the hadopelagic zone (deeper than 6,000 m), the relative abundance of JH545 decreased with increasing depth (Fig. 6b). Given the overall decline of cell numbers with increasing water depth^56–58^, the JH545 population is likely found throughout the ocean water column, reaching its highest concentration in the epipelagic zone but increasing in relative abundance in the dark ocean. This inference is also consistent with recent studies reporting higher SAR202 cell abundance in the euphotic zone compared with the aphotic zone in the Atlantic Ocean and Fram Strait as measured by FISH-based cell counting^12, 59^. The high absolute abundances of SAR202 group I members observed in the euphotic zones seemingly conflict with the observations of growth inhibition by light-dark cycles and death in continuous light of the cultured strains upon exposure to broad spectrum white light (Fig. 2b–c). To reconcile these observations, we hypothesize that JH545 uses its two copies of heliorhodopsin to regulate functions that are negatively impacted by light, mitigating its susceptibility to inhibition^34^, and we note that the action spectrum for light inhibition has not been determined and the cells could be sensitive to frequencies that are normally absorbed in the water column. Regardless, our findings indicate that the challenge of culturing these cells might in part be explained by their sensitivity to light.

**Fig. 6.**
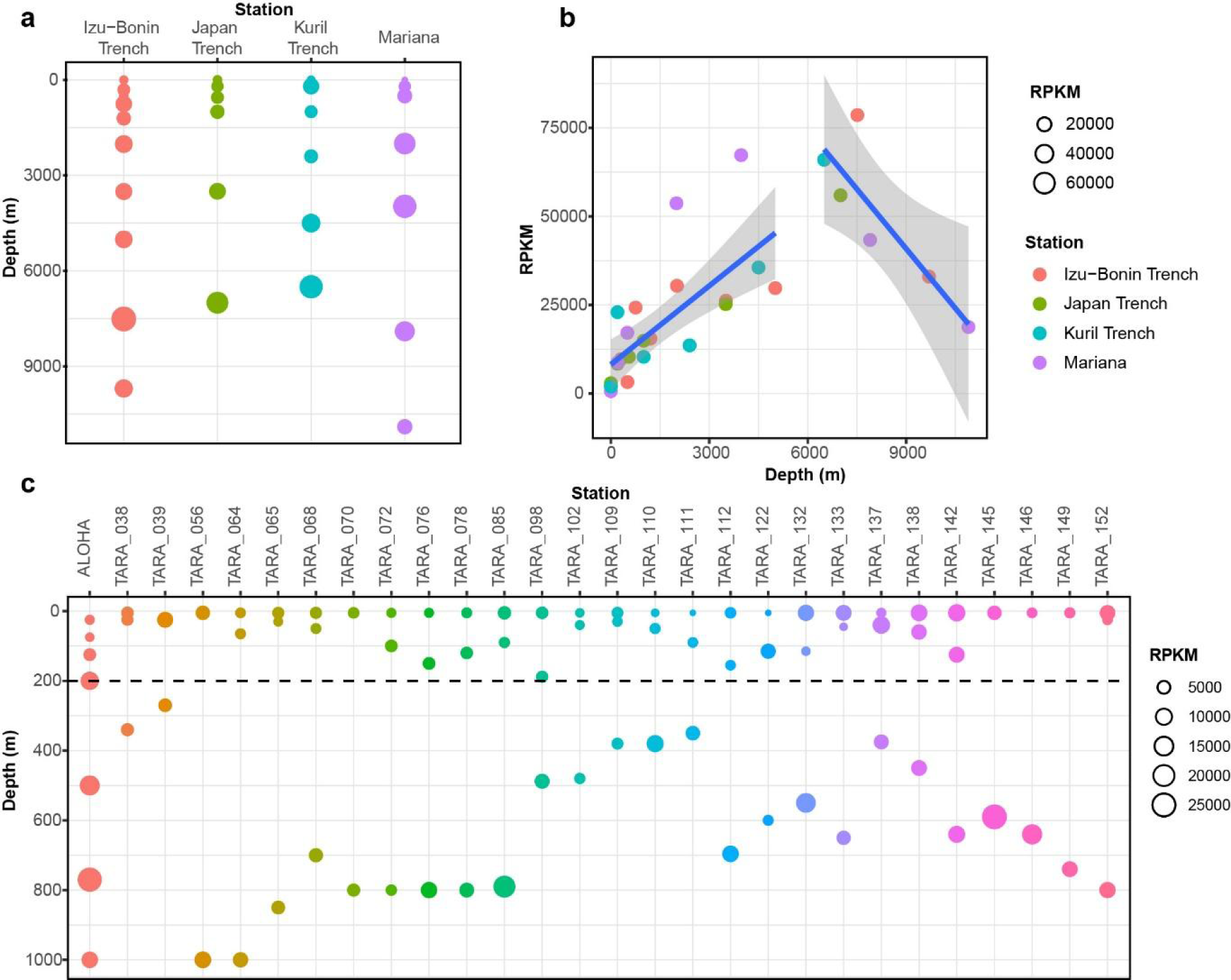
Metagenome fragment recruitment to the JH545 genome. **a,c,** RPKM values (multiplied by 10^6^) are plotted for various stations and depths from the four marine trenches with water depths of more than 6,000 m (**a**) and station ALOHA and *TARA* Oceans stations (**c**). For *TARA* Oceans, only the stations with samples from water depths of both less than 100 m and more than 250 m were used. The dotted horizontal line in (**c**) indicates a depth of 200 m, the boundary between epipelagic (euphotic) and mesopelagic zones. **b,** The data used in (**a**) are plotted irrespective of trenches to show the changes along the water depth more clearly.

We propose that the vertical distribution SAR202 group I members can be reconciled with their metabolic features that we report here. The compounds that enhanced the growth of JH545 (*e.g.*, fucose and rhamnose) and related compounds that we predict JH545 and SAR202 group I bacteria might also utilize are found in the surface ocean largely as monomers in polysaccharides produced by phytoplankton^51^. The JH545 genome had a limited repertoire of glycoside hydrolases (GHs) and lacked representative GH families annotated as fucosidase and rhamnosidase (*e.g.*, GH29, GH95, GH141, GH78, and GH106)^41, 50, 60^. SAR202 group I may scavenge monomeric compounds and their metabolic products that diffuse into the water column during degradation of polysaccharides by other taxa (e.g., *Bacteroidia*, *Gammaproteobacteria*, and *Verrucomicrobiota*), many of which are capable of much more rapid growth and would have a competitive advantage as specialists when polysaccharides are available. In the dark ocean, where polysaccharides are depleted, however, paralogous gene expansions may provide group I with a competitive advantage by allowing them to use a diverse range of sugars, sugar acids, and related compounds, leading to the observed increase in relative abundance. Piezotolerance of SAR202 cells would also give them a competitive advantage in the dark ocean^61^.

Evolutionary diversifications of paralogous enzymes in SAR202 cells have been proposed to benefit these cells by expanding the range of organic compounds they can metabolize^8, 12^. To explain the non-phosphorylative pathways for sugar oxidation we report, and the possibility of a much larger set of parallel pathways in the same cells, we hypothesize that SAR202 group I cells have evolved to exploit relatively rare carbohydrate compounds that are not harvested by taxa specializing in more common carbohydrate types. Carbohydrates are structurally complex because of the chirality of monomers, the diversity of the linkages they form, and modifications such as *O*-methylation and sulfation. In this scenario, SAR202 exploits a niche, harvesting a class of compounds that are recalcitrant because of their low abundance and structural diversity. This combination of features, *i.e.*, high molecular complexity and low concentrations of individual molecular species, is associated with the molecular diversity hypothesis^62^, a leading explanation for the sequestration of ocean carbon in dissolved molecules with millennial turnover times. Our findings are not contrary to this hypothesis; they instead suggest a class of compounds that would accumulate for the same reasons if a particular cell type, such as SAR202 group I, had not evolved a unique mechanism to harvest them.

## Conclusion

We describe the first isolates of the abundant and ubiquitous marine bacterial SAR202 clade, a monophyletic superorder within the class *Dehalococcoidia* of the phylum *Chloroflexota*. The SAR202 strains grew very slowly, and their growth was further inhibited by exposure to light; these properties might explain why they have not been cultivated previously. We show that these cells contain paralog expansions and show, for the first time, that these paralogs can be arranged into non-phosphorylative pathways for the catabolism of sugars and their lactone and acid forms. We show that fucose and rhamnose, predicted substrates of these pathways, experimentally enhance the growth of the SAR202 isolate. This is the first exploration of the physiology of a superorder of bacteria that have been implicated in the oxidation of a variety of forms of semi-labile organic carbon, and it seems likely that further studies of the diverse cells in this clade will contribute to a more mechanistic understanding of the ocean carbon cycle.

## Methods

### Sample collection and cultivation

Seawater samples used for high-throughput culturing (HTC) based on dilution-to-extinction and amplicon analysis were collected from a depth of 10 m at two stations (GR1 and GR3) in the West Sea of Korea (Yellow Sea) in October 2017 (Extended Data Fig. 1a). Physicochemical properties of the water samples are presented in Extended Data Fig. 1b. The total prokaryotic number was determined by counting 4′,6-diamidino-2-phenylindole (DAPI)-stained cells using an epifluorescence microscope (Nikon 80i, Nikon) after filtration using a 0.2-μm pore-sized polycarbonate membrane filter (Millipore). Culture media for HTC (LNHM) was prepared using a seawater sample collected from the East Sea (depth, 10 m) in 2016. In brief, the seawater sample was filtered using a 0.2-μm pore-sized polyethersulfone membrane filter (Pall), autoclaved (1.5 h), sparged with CO_2_ (8 h), aerated (24 h), and amended with carbon sources, macronutrients (nitrogen and phosphorus sources), trace metals, vitamins, and amino acids (Extended Data Table 1). The seawater samples were then diluted with culture media to a concentration of 5 cells mL^-1^ and dispensed (1 mL per well) into 48-well microplates (BD Falcon). The plates were incubated at 20 °C for 1 month in dark or under LED light (Philips; correlated color temperature, 3000 K; light intensity, ∼155 μmol photons m^⁻2^ s^⁻1^) with a light/dark cycle of 14:10 h. The microbial growth in each well was screened by flow cytometry (GUAVA EasyCyte Plus flow cytometer, Millipore) after staining with SYBR Green I (Life Technologies) and recorded as growth-positive when more than 5.0 × 10^4^ cells mL^-1^ were detected. Cultures from growth-positive wells were used for further analyses and stored as glycerol stock (10%, v/v) at −80 °C.

### Phylogenetic analysis and classification of 16S rRNA sequences

Phylogenetic analyses of growth-positive cultures were based on PCR amplification and sequencing of 16S rRNA genes. DNA templates for PCR were prepared from growth-positive cultures using the InstaGene^TM^ Matrix (Bio-Rad) according to the manufacturer’s instructions. The amplification of 16S rRNA genes were performed by PCR using 27F and 1492R primers, followed by Sanger sequencing with 800R and 518F primers (Macrogen Inc., Korea). Taxonomic classification of 610 strains obtained from the HTC experiments was carried out using the “classify.seqs” command of the Mothur software package (v1.39.5)^63^ using the SILVA database SSURef NR99 (release 132)^64^ as a reference. For more refined taxonomic and phylogenetic analysis of the SAR202 isolates, the 16S rRNA gene sequences were aligned using the SINA online aligner (v1.2.11, http://www.arb-silva.de/aligner)65, imported into ARB program^66^, and inserted using the ARB parsimony into the guide tree of SILVA database SSURef NR99 (release 132)^64^. After manual curation, the aligned sequences of the isolates and their phylogenetic relatives were exported with “ssuref:bacteria” filter. Maximum-likelihood phylogenetic trees were constructed using RAxML^67^ (v8.2.12) with GTRGAMMA method including 100 bootstrap replicates and visualized using the MEGA software (v7.0)^68^.

### Microbial community analysis

Two liters of each seawater sample (GR1 and GR3) were filtered through a 0.2-μm pore-size polyethersulfone membrane filter (Supor, Pall). DNA was extracted directly from the membrane filters using DNeasy PowerWater Kit (Qiagen) according to the manufacturer’s instructions. The V4-V5 regions of 16S rRNA genes were amplified using fusion primers that were designed based on universal primers, 518F and 926R^69^. The pooled PCR products were sequenced on Illumina MiSeq platform (300-bp, paired-end; Chunlab Inc.). The analyses of the 16S rRNA gene amplicon sequences were performed using QIIME2^70^ after primer trimming with cutadapt (v2.7)^71^.

### Culture experiments

The SAR202 strains were grown and maintained using LNHM5x at 20 °C in dark. Growth experiments were performed using LNHM, LNHM5x, and ASW5x. The detailed recipes of the media are presented in Extended Data Table 1. Bacterial growth was monitored by flow cytometry, and the purity and identity of cultures were regularly determined by sequencing the amplified 16S rRNA genes and by microscopic examination.

The growth of strain JH545 was monitored at temperatures ranging from 4 ℃ to 37 °C in dark. Growth substrate tests were performed using ʟ-rhamnose, ʟ-rhamnono-1,4-lactone, ʟ-rhamnonate, ʟ-fucose, ʟ-fucono-1,4-lactone, ʟ-fuconate, and ascorbate. All tested compounds were purchased from Sigma-Aldrich. These chemicals were added to ASW5x media at a final concentration of 250 μM. Cultures were incubated in dark at 20 ℃.

For the initial examination of the effect of light exposure on SAR202 growth (Fig. 2b), cells were inoculated in LNHM at an initial cell density of ∼1.0 × 10^4^ cells mL^-1^ and incubated in a chamber equipped with LED (3000 K; Phillips), with a light/dark cycle of 14:10 h. Light intensity was ∼155 μmol photons m^-2^ s^-1^. To test the effect of light intensity on SAR202 growth (Fig. 2c), a custom-made equipment with a LED-array (3000 K) and a light dimmer was utilized, and the light intensities were set at ∼45 (weak light), ∼89 (medium light), and ∼134 (strong light) μmol photons m^-2^s^-1^. During the test, the lights continuously remained on. All experiments included control samples that were constantly kept in dark by wrapping the tubes or flasks with aluminum foil. Light intensity was measured using a digital light meter (TES-1335; TES).

### Morphological characterization

Cell morphology was observed by transmission electron microscopy (TEM; CM200, Philips) and scanning electron microscopy (SEM; S-4300 and S-4300SE, Hitachi). To prepare samples for TEM, 20 mL of culture prefixed with 2.5% glutaraldehyde were filtered using a 0.2-μm pore-size polycarbonate membrane on which formvar/carbon-coated copper grids were placed, followed by staining of the grids with uranyl acetate (2%). For thin-section TEM, centrifuged cells from 800 mL of culture were subjected to primary fixation with Karnovsky’s solution (2% paraformaldehyde, 2.5% glutaraldehyde), post-fixation with 2% OsO_4_, en-bloc staining with 0.5% uranyl acetate, sequential dehydration with 30, 50, 70, 80, 90, and 100% ethanol, and embedding with resin (EMBed-812, Electron Microscopic Science). Finally, sectioning was performed using a diamond knife. Thin sections were stained with uranyl acetate (2%). For SEM analysis, 100 mL of culture was concentrated by centrifugation, fixed with 2.5% glutaraldehyde, post-fixed with 1% OsO_4_, dehydrated with 30, 50, 70, 80, 90, and 100% ethanol, and chemically dried using hexamethyldisilazane (Sigma-Aldrich). Treated samples were gently mounted on a cover glass and coated with a thin layer of carbon.

### Genome sequencing, assembly, and analyses

Genomic DNA of the SAR202 strains was extracted from cell pellets obtained by centrifugation of liquid cultures (∼1 L), using DNeasy Blood & Tissue Kit (Qiagen), according to manufacturer’s instructions. The genomic DNA of strains JH545 and JH1073 was used for the construction of the 20-kb SMRTbell library, which was sequenced on the PacBio RS II platform (Pacific Biosciences). *De novo* assembly of raw sequencing reads was carried out by the RS_HGAP_Assembly.2 protocol of SMRT Analysis (v2.3.0), resulting in a single contig. The contig was circularized using Circlator (v1.5.5)^72^ and polished using the RS_Resequencing.1 protocol of SMRT Analysis to obtain the final error-corrected genome sequence. Genome sequencing of strains JH702 and JH639 was performed on the Illumina HiSeq platform (2 × 150 bp). Raw reads were trimmed using BBDuk with the following options: ktrim=r k=23 mink=11 hdist=1 tpe tbo ftm=5 qtrim=rl trimq=10 minlen=100. Assembly of Illumina sequencing data was performed using SPAdes v3.11.1 in a multi-cell mode with read error and mismatch correction^73^.

The genome sequences were submitted to the IMG-ER system for annotation. Prokka (v1.12)^74^ was also used for annotation. The predicted protein sequences were analyzed using BlastKOALA^75^ and KofamKOALA^76^ for metabolic pathway reconstruction based on KEGG Orthologs (KOs). Annotation by eggnog-mapper (v2.0.1)^77^ and hmmsearch (v3.3) against protein databases such as Pfam were also performed for more accurate and detailed functional annotation. Analysis of CAZymes was performed using dbCAN2^78^. A map of metabolic pathways was created using BioRender (https://biorender.com). ANIb values between genomes were calculated using JSpeciesWS^79^. Genomic islands were predicted using IslandViewer 4^80^. Genome comparison was visualized by BLAST Ring Image Generator (BRIG)^81^.

The SAR202 genome sequences of the present study and a previous study^12^ were classified using GTDB-Tk, which indicated that the genomes belonged to ∼10 different orders according to the GTDB. Species cluster-representative genomes of these orders were used to construct a phylogenomic tree of the SAR202 clade. We used UBCG pipeline^82^ to obtain a concatenated alignment of core genes. A maximum-likelihood tree was constructed using RAxML (v8.2.12)^67^, with a PROTGAMMAAUTO option including 100 bootstrap iterations.

### Analysis of the HeR

Genomic regions around the HeR genes of the JH545 genome were visualized using Easyfig^83^. Multiple amino acid alignment of HeRs from JH545 and the first-characterized HeR (48C12) was performed using ClustalW^84^. The Clustal X colour scheme of the Jalview (v2.11.1.3)^85^ was applied to visualize the alignment.

### Analysis of COG4948 proteins

A total of 80 proteins were assigned to COG4948 by the IMG/ER annotation of the JH545 genome, which was also verified by the Conserved Domain search^86^ (v3.18) at NCBI. A search against Pfam-A database (by Pfam_Scan.pl; both available at https://ftp.ebi.ac.uk/pub/databases/Pfam/) showed that 74 COG4948 proteins had both PF02746 and PF13378 domains. The remaining six proteins had either PF13378 (four proteins) or PF02746 (two proteins) domains. The two Pfam domains were found only in 80 COG4948 proteins. Based on these results showing a correspondence between COG4948 and the two Pfam domains, we decided to use the two Pfam domains for approximate calculation of the proportion of COG4948 proteins in 47,894 species cluster-representative genomes of the GTDB (R202). The genomic faa files (gtdb_proteins_aa_reps_r202.tar.gz) were downloaded from the GTDB repository (https://data.gtdb.ecogenomic.org/) and searched using hmmsearch (v3.3) against the hmm files of the two Pfam domains, with “cut_tc” option. The approximate proportion of COG4948 proteins in each genome was calculated by dividing the number of hmmsearch hits by twice the number of CDS. The results were visualized using the R package ‘tidyverse’.

The 80 COG4948 protein sequences of JH545 were submitted to the Enzyme Similarity Tool (EFI-EST; https://efi.igb.illinois.edu/efi-est/)42 to generate an SSN. A total of 77,142 proteins in the UniProt database (v2020_02) that had the two Pfam domains were included in the SSN, which was constructed using BLAST with default options. The obtained SSN was explored using the Cytoscape (v3.7.2)^87^. After the application of a cutoff threshold of alignment score (larger than or equal to 150), clusters that included the 80 COG4948 proteins of the strain JH545 were retained for visualization.

Reconstruction of non-phosphorylative fucose and rhamnose degradation pathways is based on a KEGG pathway map (fructose and mannose metabolism; map00051), MetaCyc pathways (L-fucose degradation II and L-rhamnose degradation II/III), and several publications^46, 47, 52–55^. When necessary, proteins characterized in past reports were explored in the UniProt database or analyzed by CD-search at NCBI to ascertain their COG assignments.

### Metagenome fragment recruitment

Metagenomes from several marine trenches, station ALOHA, and some *Tara* Oceans stations were downloaded from ENA, quality-trimmed using BBduk (v38.86)^88^, subsampled to 1 million reads using Seqtk (https://github.com/lh3/seqtk), and used for BlastN against the JH545 genome. Ribosomal RNA and tRNA genes of the JH545 genome were masked before the analyses. The number of BlastN hits (*i.e.*, metagenome reads) that satisfied the cutoff values of identity (at least 90%) and alignment length (at least 50 bp) were counted for each metagenome and used for the calculation of RPKM values. The list of metagenome samples is provided in Supplementary Table 1.

## Data availability

The 16S rRNA gene sequences of 24 isolates in SAR202 group I are available under the GenBank accession numbers OQ689977-OQ690000. The whole genome sequences have been deposited in GenBank with accession numbers CP046146 (JH545), CP046147 (JH1073), WMBD00000000 (JH639), and WMBE00000000 (JH702), and also available at IMG/M database with genome IDs 2901382945 (JH545), 2917498938 (JH1073), 2892960865 (JH639), and 2892963810 (JH702).

## Acknowledgments

This research was supported by High Seas Bioresources Program of Korea Institute of Marine Science & Technology Promotion (KIMST) funded by the Ministry of Oceans and Fisheries (KIMST-20210646), by the Mid-Career Research Program (NRF-2022R1A2C3008502), and by the Science Research Center Grant (NRF-2018R1A5A1025077) through the National Research Foundation (NRF) funded by the Ministry of Sciences and Information and Communications Technology, Korea.

## Author contributions

J.-H.S. and J.-C.C. planned and designed the initial isolation project. Y.L. and J.-H.S. performed the experiments. Y.L., J.-H.S., and I.K. analyzed the data. Y.L, I.K, S.J.G, and J.-C.C. wrote and revised the manuscript. I.K and J.-C.C. supervised the project.

## Supplementary information

The online version contains supplementary material available at the website.

---

**Extended Data Fig. 1.**
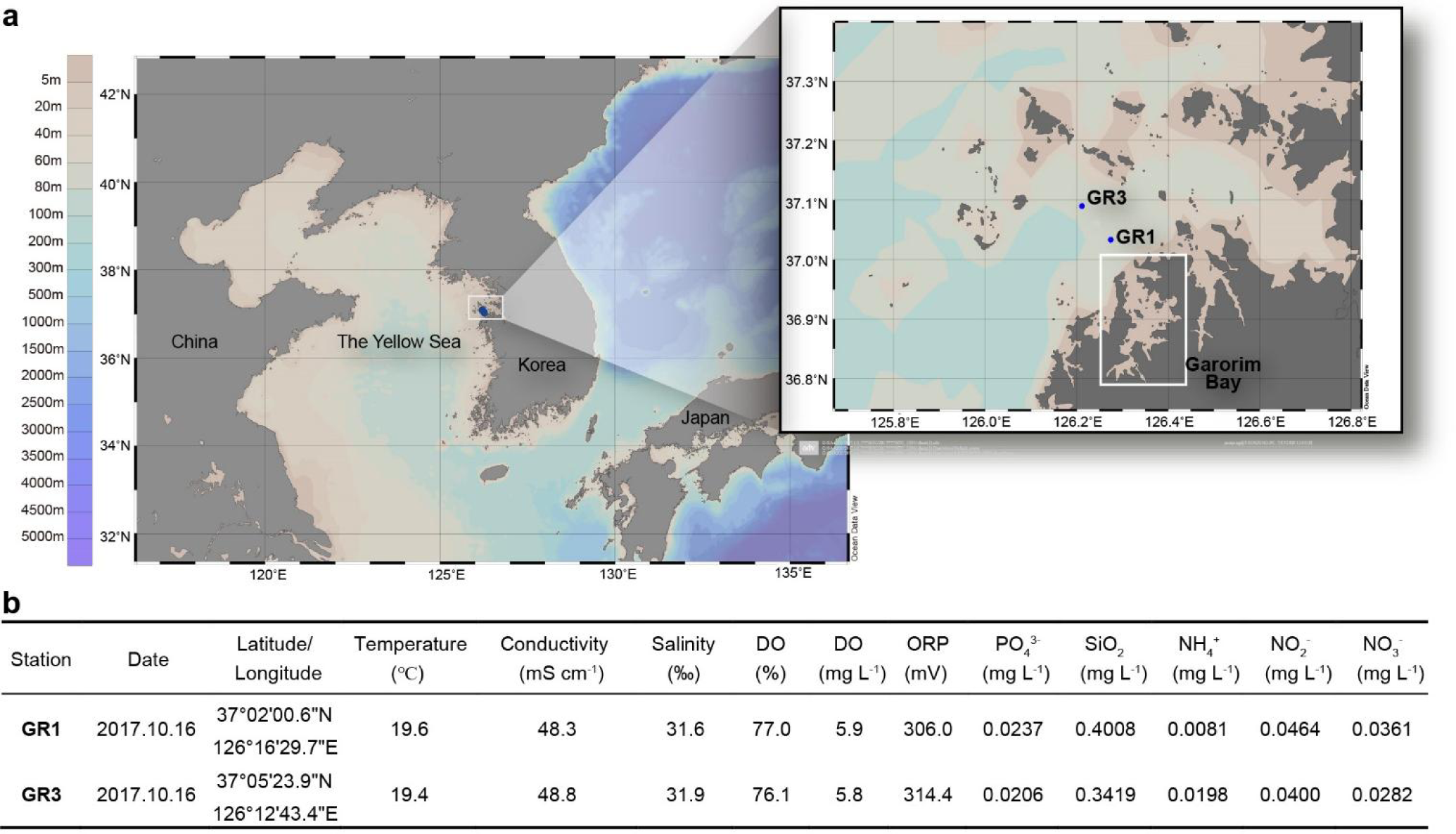
Map of the sampling stations and physicochemical parameters of the seawater samples used for HTC and amplicon sequencing. **a,** Map of the sampling stations. The two blue dots indicate the sampling stations, GR1 and GR3, located outside of the Garorim Bay in the Yellow Sea. The map was created by Ocean Data View program. **b,** Physicochemical parameters of the seawater samples used for HTC and amplicon sequencing. Temperature, conductivity, salinity, dissolved oxygen (DO), and oxidation reduction potential (ORP) were measured using a portable instrument (YSI Model 556, YSI Incorporated). Concentrations of inorganic nutrients were determined using a QuAAtro microflow analyzer (SEAL Analytical).

**Extended Data Fig. 2.**
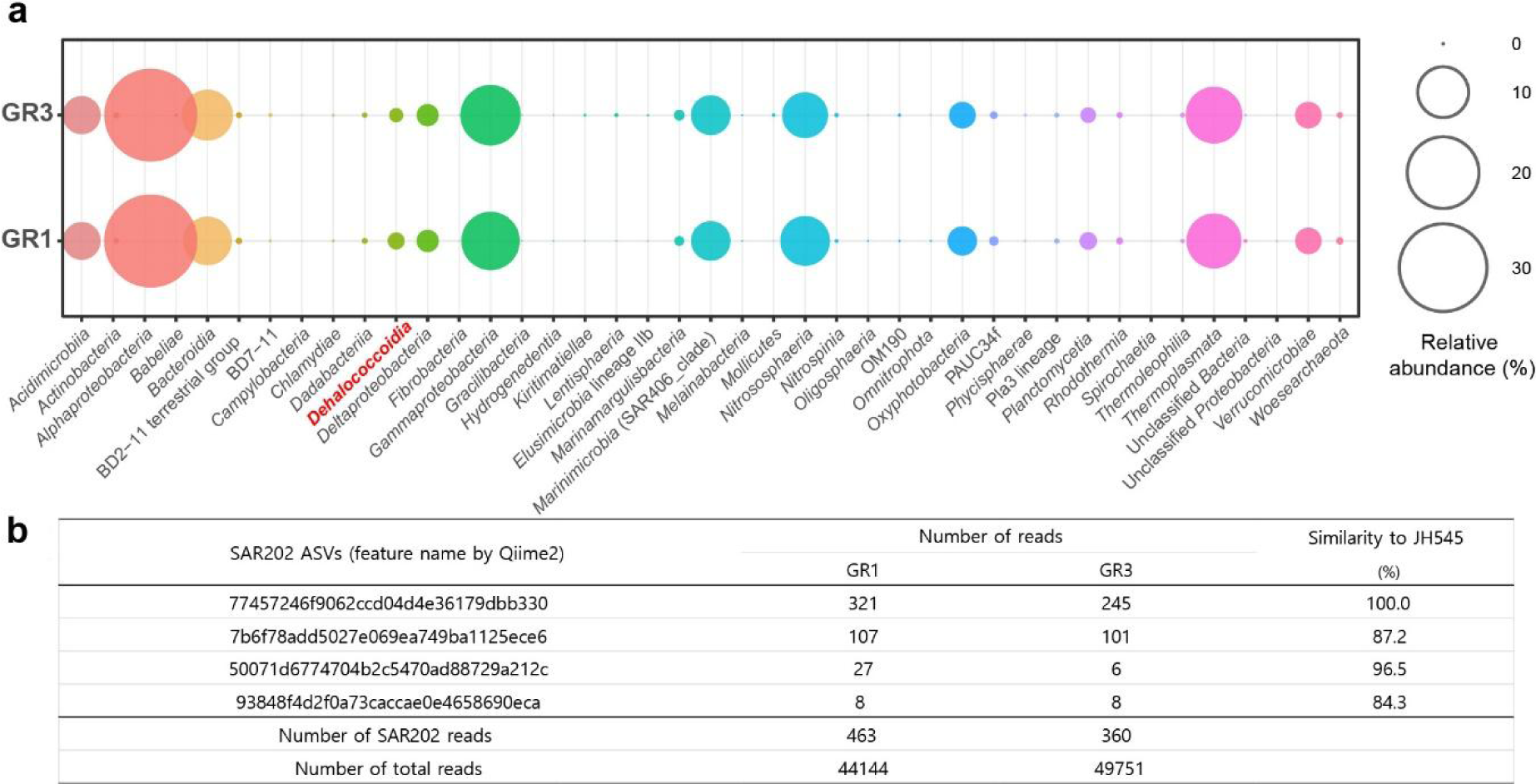
Prokaryotic community composition of the seawater samples used for the HTC experiment, collected at sampling stations GR1 and GR3. The sequencing data of 16S rRNA gene amplicon were analyzed by DADA2 plugin of QIIME2. Taxonomic classification of ASVs was based on SILVA SSU database (release 138). a, Relative abundance of prokaryotic groups at class level. Most of the reads classified into *Dehalococcoidia* (indicated in red) belonged to the SAR202 clade. b, The number of sequencing reads assigned to the four SAR202 ASVs in each seawater sample. Sequence similarities between JH545 and the ASVs are also indicated.

**Extended Data Fig. 3.**
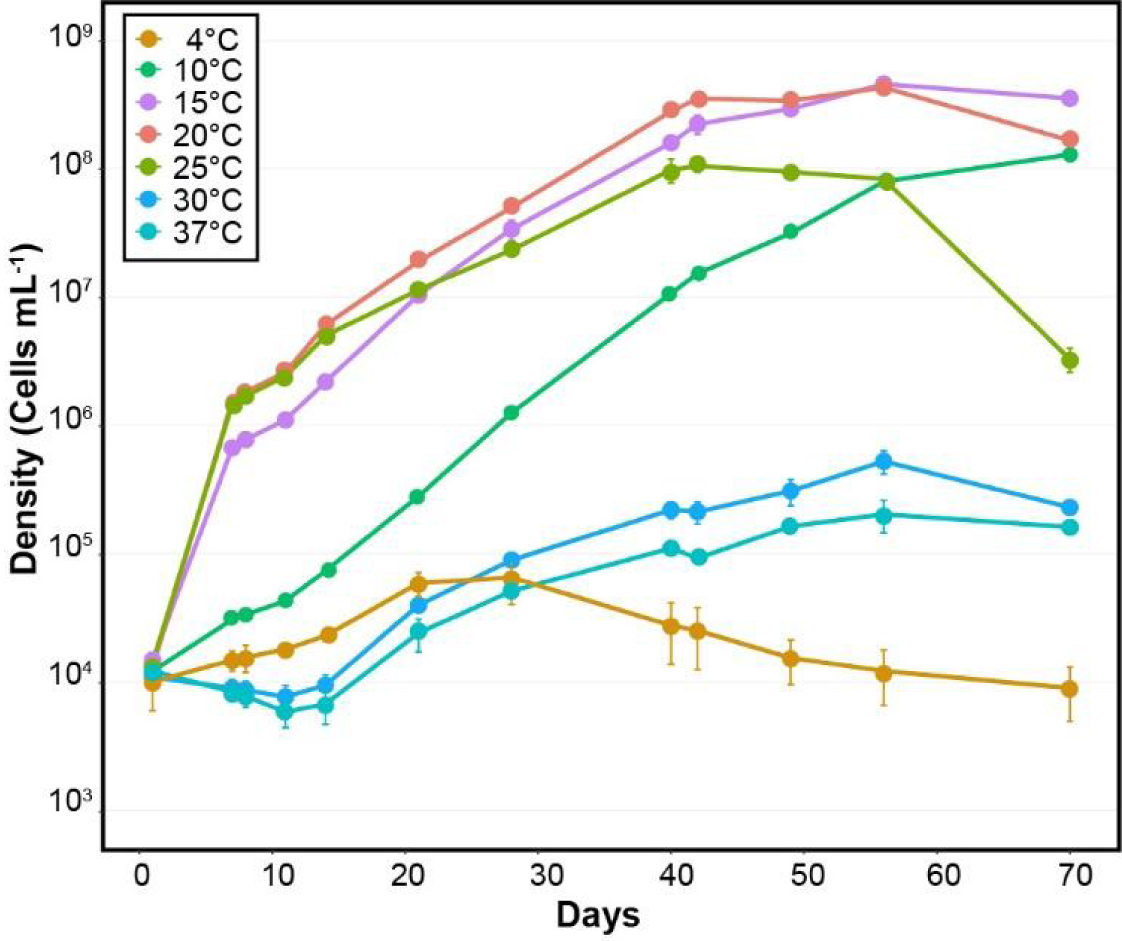
Growth curves of strain JH545 at various temperatures. Cultures were incubated in LNHM in dark at temperatures ranging between 4 ℃ and 37 ℃. Experiments were performed in triplicates. Error bars indicate standard deviation. Note that the error bars are hidden when they are shorter than the size of the symbols.

**Extended Data Fig. 4.**
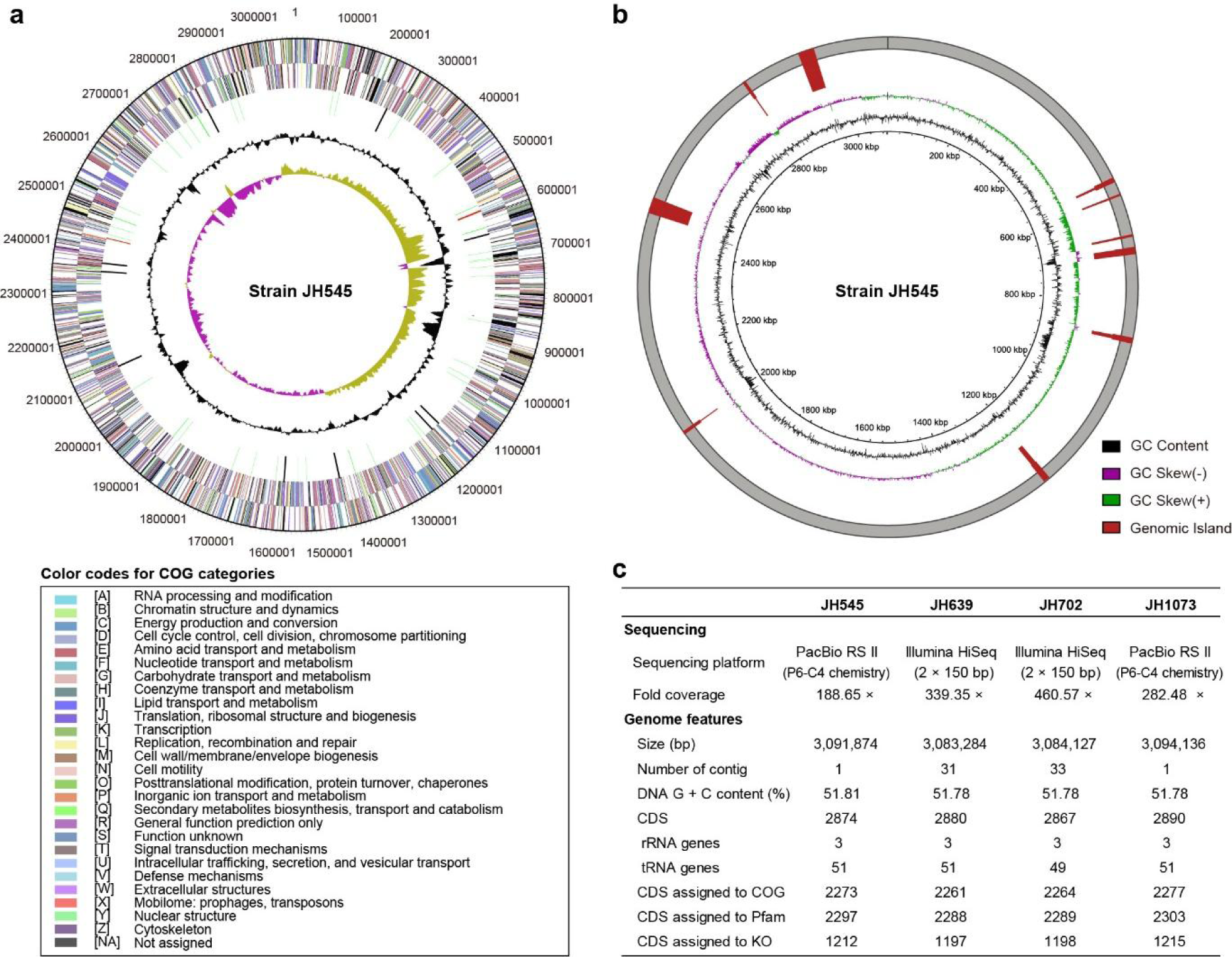
Features of the four SAR202 genomes. **a,** Genome map of strain JH545 as drawn by IMG. From outside to the center: Genes on the forward strand (colored by COG categories), genes on the reverse strand (colored by COG categories), RNA genes (tRNAs in green, rRNAs in red, other RNAs in black), GC content, and GC skew. **b,** Genomic islands of the JH545 genome predicted by IslandViewer 4. The two inner rings showing GC content and GC skew were drawn by BRIG (BLAST ring image generator). The outermost ring shows the positions of genomic islands. Note that only the integrated results are shown. **c,** Genome statistics of the four SAR202 strains. Annotation was performed by the IMG-ER pipeline.

**Extended Data Fig. 5.**
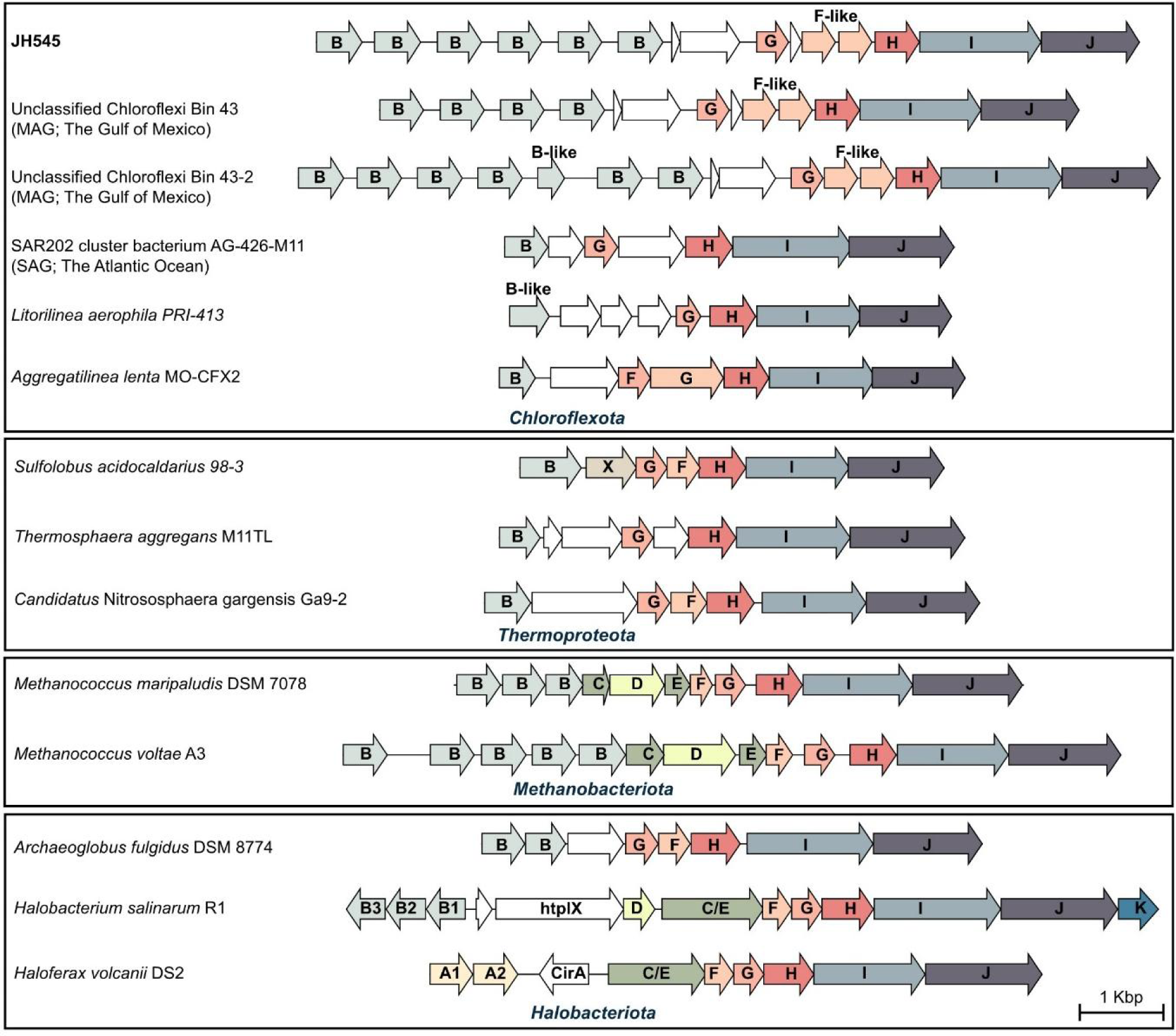
Map of archaella gene clusters found in the JH545 genome, three SAR202 MAG/SAGs, two *Chloroflexota* isolates, and several archaeal genomes. Gene names are shown inside the arrows representing genes. Names of archaella genes are shown with a single capital letter that corresponds to the letter “X” in the gene name “flaX”, *i.e.*, the last letter of the gene name. Homologous genes are shown in the same color. Genes of unknown function are depicted in white. The flaB-like and flaF-like genes of the SAR202 genomes show sequence similarity to the respective archaeal homologs but were not annotated clearly. Genomes are grouped by their phylum-level affiliations according to the GTDB. Easyfig was used to generate the figure.

**Extended Data Fig. 6.**
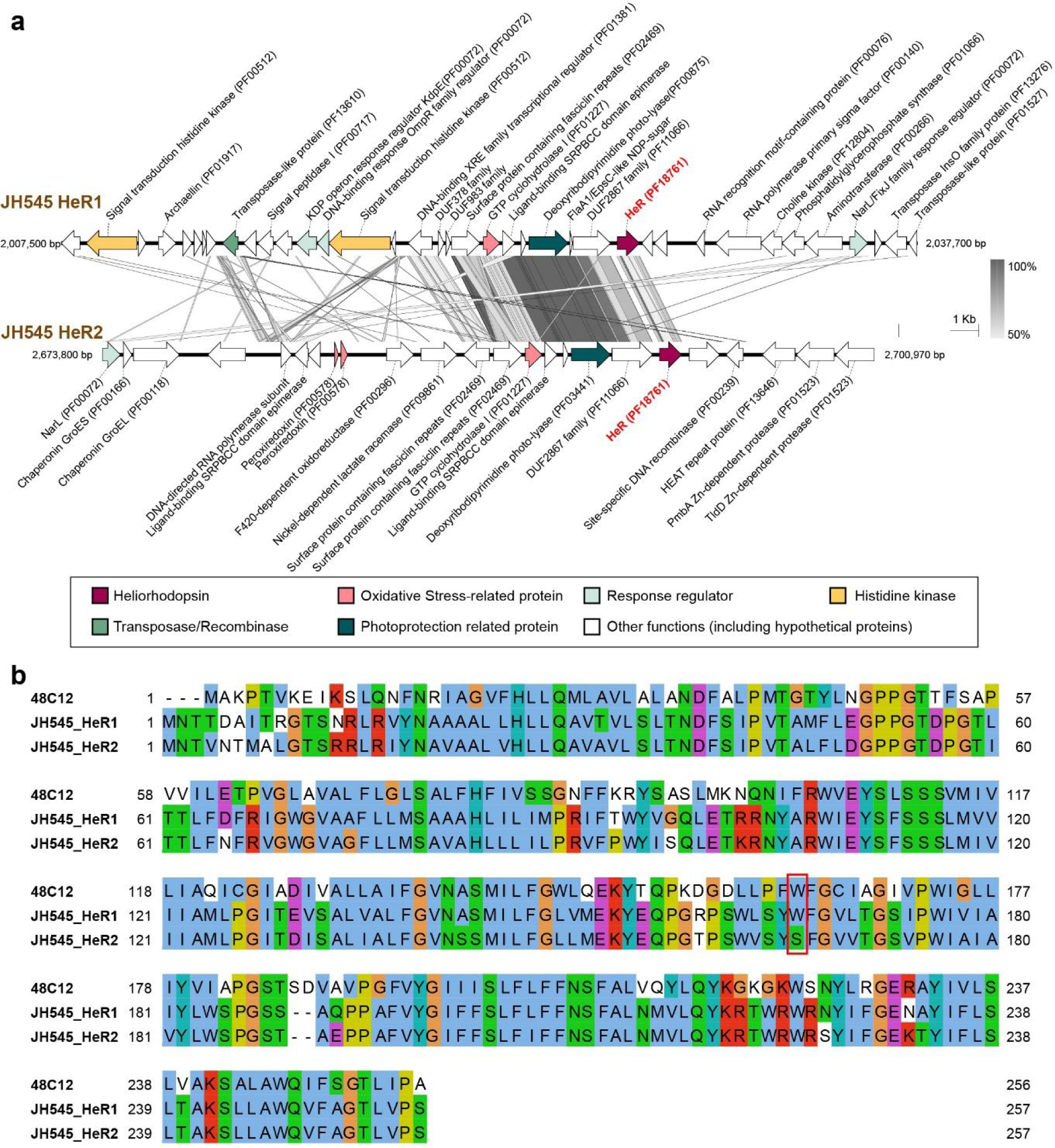
Heliorhodopsin (HeR) in the JH454 genome. **a,** Genomic context around the two heliorhodopsin genes of the JH545 genome. Genes putatively assigned to several functions are indicated by colors following the legend at the bottom. Grey boxes between the two genomic regions indicate nucleotide-level similarity following the scale at middle right. This map was drawn using Easyfig. **b,** Amino acid sequence alignment of HeR proteins. The two HeR proteins of JH545 and the first characterized HeR (48C12) were aligned by Clustal Omega and visualized by Jalview with Clustal X color scheme. The red box indicates the amino acid position (W163 in 48C12) where the two HeR of JH545 have different residues and replacement with alanine caused a spectral shift in 48C12.

**Extended Data Fig. 7.**
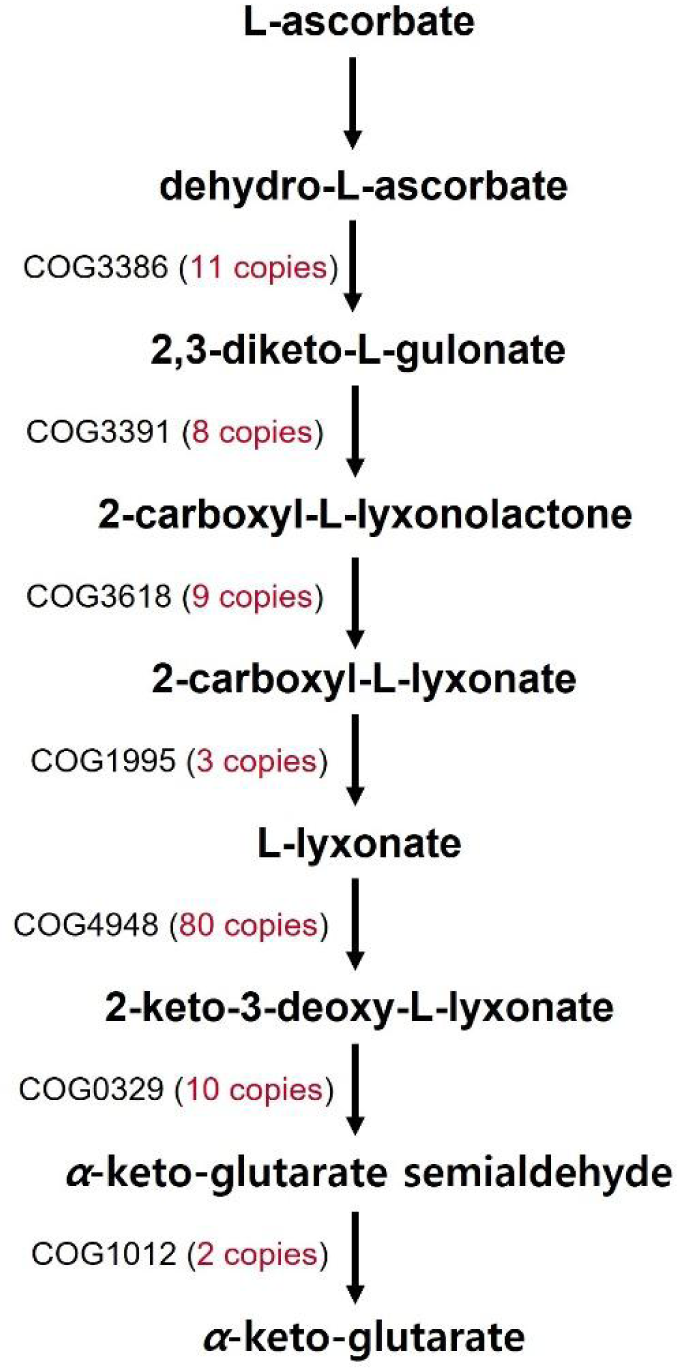
Catabolic pathways of L-ascorbate inferred from SAR202 genome annotation. This non-phosphorylative pathway is based on a KEGG pathway map (ascorbate and aldarate metabolism; map00053) and MetaCyc pathways (L-ascorbate degradation and L-lyxonate degradation). The number of proteins assigned to the COGs are indicated within parentheses next to the COG IDs.

**Extended Data Table 1.**
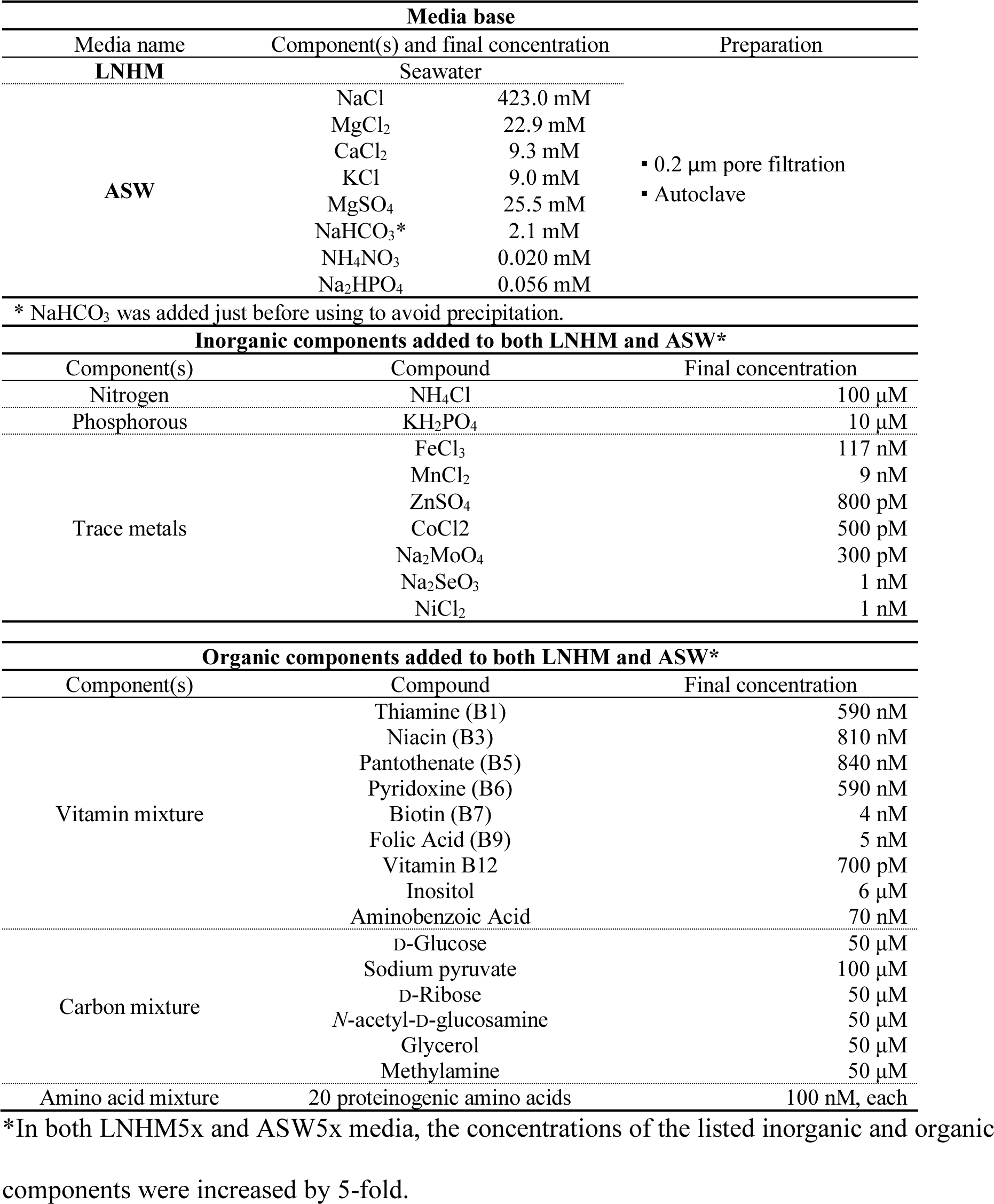
The compositions of low-nutrient heterotrophic medium (LNHM) and artificial seawater (ASW) medium used in this study.

**Extended Data Table 2.**
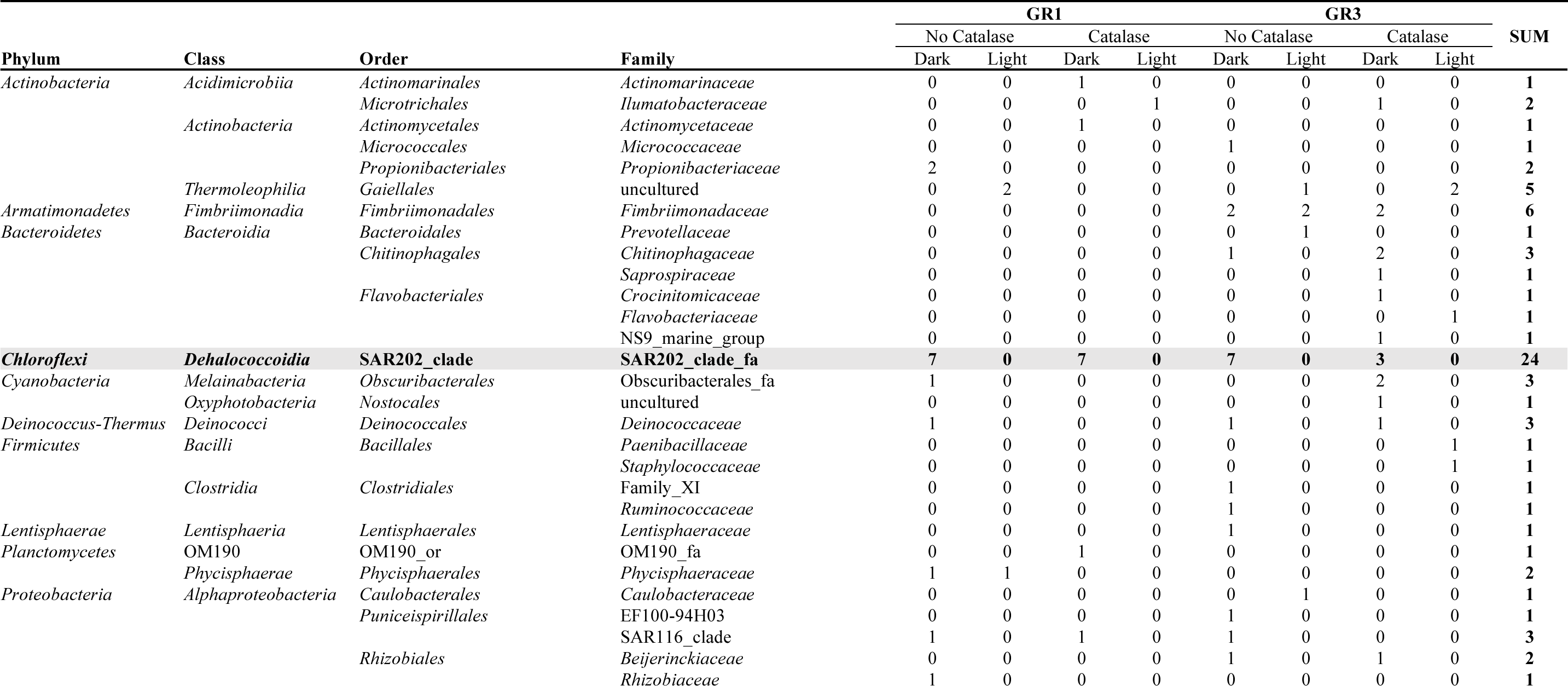
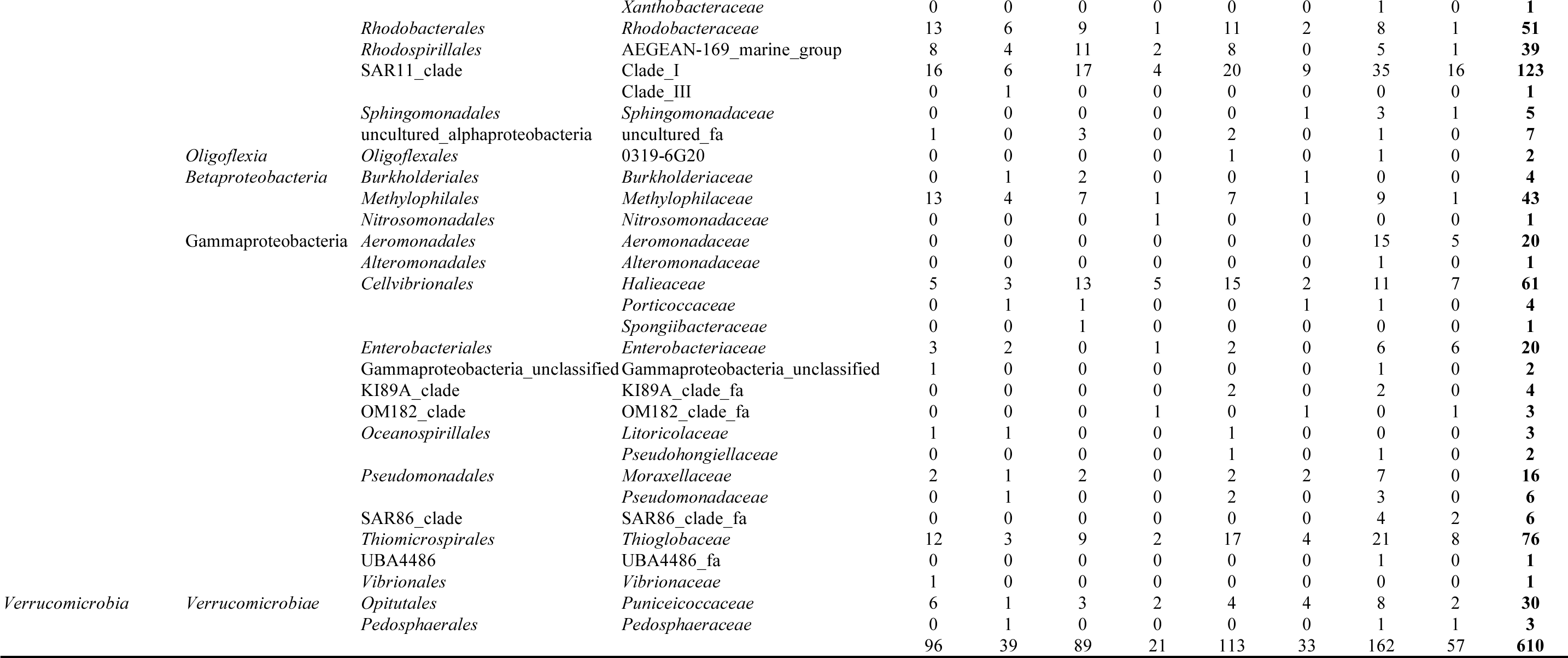
Taxonomic distribution of the 610 isolates obtained by HTC experiments. Taxonomic classification was performed using Mothur (v1.39.5; “classify.seqs” command) based on SILVA SSUREF database (release 138) and is presented at the family level. Sampling stations and culture conditions (catalase addition, dark vs. light) are also indicated. Note that “_or” and “_fa” in some order and family names are suffixes indicating the taxonomic level of the taxa.

**Extended Data Table 3.** The 30 most abundant COGs found in the SAR202 genomes obtained in this study and the number of proteins assigned

| COG IDs | Description of COG | No. of CDS |  |  |  |
| --- | --- | --- | --- | --- | --- |
|  |  | JH545 | JH639 | JH702 | JH1073 |
| COG4948 | L-alanine-DL-glutamate epimerase or related enzyme of enolase superfamily | 80 | 80 | 79 | 79 |
| COG1028 | NAD(P)-dependent dehydrogenase, short-chain alcohol dehydrogenase family | 44 | 44 | 44 | 45 |
| COG0477 | MFS family permease | 42 | 42 | 41 | 42 |
| COG0642 | Signal transduction histidine kinase | 40 | 41 | 41 | 40 |
| COG0451 | Nucleoside-diphosphate-sugar epimerase | 24 | 24 | 24 | 24 |
| COG0745 | DNA-binding response regulator, OmpR family, contains REC and winged-helix (WHTH) domain | 23 | 23 | 23 | 23 |
| COG3836 | 2-keto-3-deoxy-L-rhamnonate aldolase RhmA | 19 | 19 | 19 | 19 |
| COG2197 | DNA-binding response regulator, NarL/FixJ family, contains REC and HTH domains | 18 | 18 | 18 | 18 |
| COG3119 | Arylsulfatase A or related enzyme | 18 | 18 | 18 | 18 |
| COG0667 | Predicted oxidoreductase (related to aryl-alcohol dehydrogenase) | 17 | 17 | 17 | 17 |
| COG0673 | Predicted dehydrogenase | 17 | 17 | 17 | 17 |
| COG0111 | Phosphoglycerate dehydrogenase or related dehydrogenase | 15 | 15 | 15 | 15 |
| COG1063 | Threonine dehydrogenase or related Zn-dependent dehydrogenase | 15 | 15 | 15 | 15 |
| COG0028 | Acetolactate synthase large subunit or other thiamine pyrophosphate-requiring enzyme | 14 | 14 | 14 | 14 |
| COG1011 | FMN phosphatase YigB, HAD superfamily | 14 | 14 | 14 | 14 |
| COG0596 | Pimeloyl-ACP methyl ester carboxylesterase | 13 | 13 | 13 | 13 |
| COG1312 | D-mannonate dehydratase | 13 | 13 | 13 | 13 |
| COG1681 | Archaeallin (archaeal flagellin) | 13 | 13 | 13 | 13 |
| COG0438 | Glycosyltransferase involved in cell wall biosynthesis | 12 | 12 | 12 | 12 |
| COG0684 | Regulator of RNase E activity RraA | 12 | 12 | 12 | 12 |
| COG0500 | SAM-dependent methyltransferase | 11 | 11 | 11 | 11 |
| COG2055 | Malate/lactate/ureidoglycolate dehydrogenase, LDH2 family | 11 | 11 | 11 | 11 |
| COG2226 | Ubiquinone/menaquinone biosynthesis C-methylase UbiE | 11 | 11 | 11 | 11 |
| COG2801 | Transposase InsO and inactivated derivatives | 11 | 11 | 2 | 3 |
| COG2963 | Transposase and inactivated derivatives | 11 | 11 | 3 | 3 |
| COG3386 | Sugar lactone lactonase YvrE | 11 | 11 | 11 | 11 |
| COG0329 | Dihydrodipicolinate synthase/N-acetylneuraminate lyase | 10 | 10 | 10 | 10 |
| COG0784 | Dihydrodipicolinate synthase/N-acetylneuraminate lyase | 9 | 9 | 9 | 9 |
| COG2010 | Cytochrome c, mono- and diheme variants | 9 | 9 | 9 | 9 |
| COG3618 | Predicted metal-dependent hydrolase, TIM-barrel fold | 9 | 9 | 9 | 9 |
| SUM of all |  | 2273 | 2261 | 2264 | 2277 |

