## Supplementary Information for "Cultivation of SAR202 Bacteria from the Ocean"

SUPPLEMENTARY FIGURES ........ 1
SUPPELMENTARY TABLES ........ 2
Supplementary Table. 1 ........ 2
Supplementary Table. 2 ........ 4
SUPPLEMENTARY TEXT ........ 5
Genome-based phylogeny of the SAR202 clade ........ 5
Proposal of taxonomic ranks of *Candidatus* Lucifugimonas marina ........ 6
Description of "*Candidatus* Lucifugimonas" gen. nov. ........ 6
Description of "*Candidatus* Lucifugimonas" marina sp. nov. . ........ 6

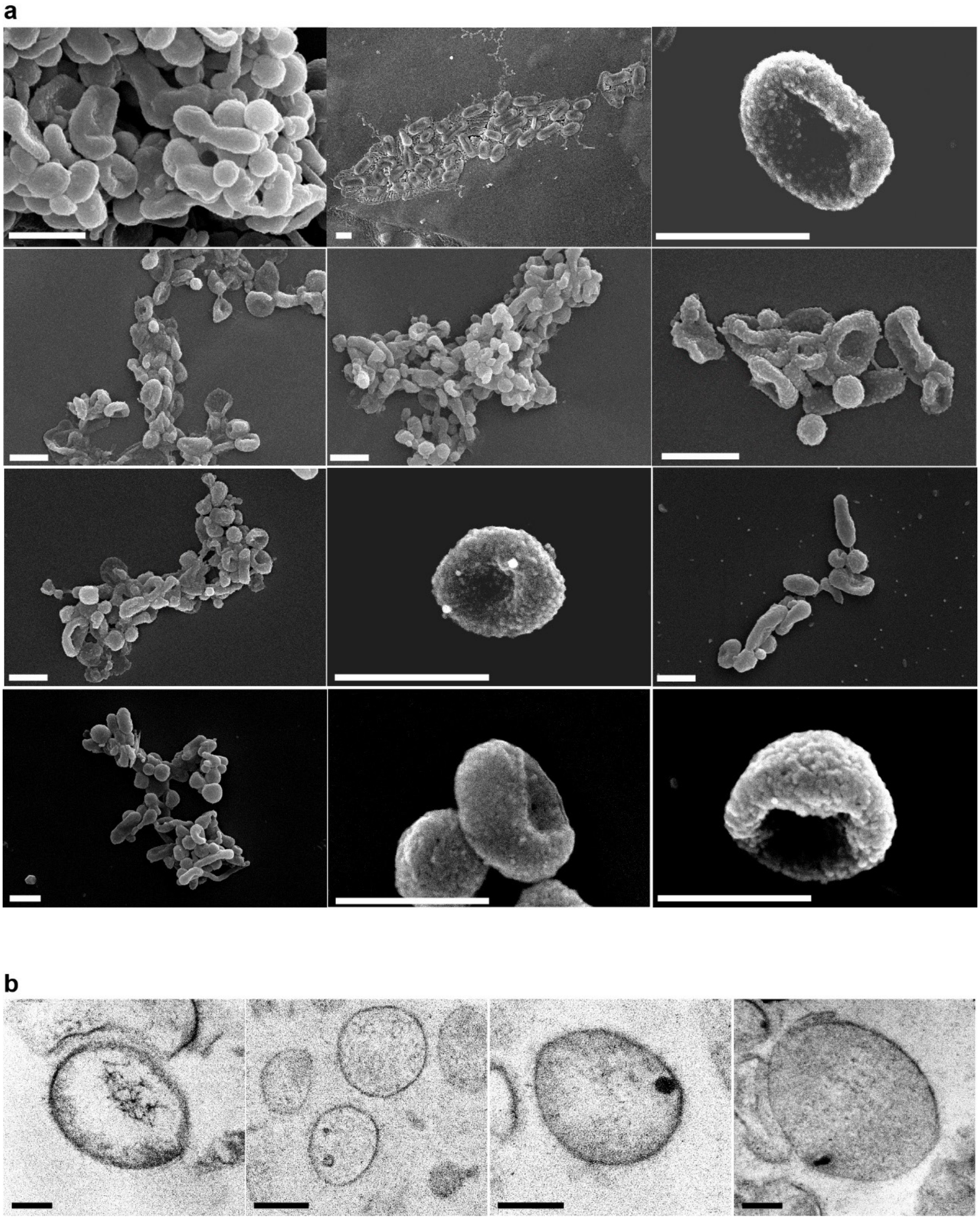

**Supplementary Fig. 1 | Morphology of strain JH545 cells observed by SEM (a) and thin-section TEM (b).** Scale bars represent 1 μm and 200 nm in panels (a) and (b), respectively.

**Supplementary Table 1 | Metagenome samples used in fragment recruitment.**

| Station | Depth (m) | Latitude | Longitude | Accession |
|---|---|---|---|---|
| TARA_038 | 5 | 19.0393 | 64.4913 | ERR599102, ERR599158 |
| TARA_038 | 25 | 19.0284 | 64.5126 | ERR598949, ERR599082 |
| TARA_038 | 340 | 19.0351 | 64.5638 | ERR599109, ERR599167 |
| TARA_039 | 25 | 18.5839 | 66.4727 | ERR599145 |
| TARA_039 | 270 | 18.7341 | 66.3896 | ERR599037, ERR599172 |
| TARA_056 | 5 | -15.3424 | 43.2965 | ERR599057 |
| TARA_056 | 1000 | -15.3379 | 43.2948 | ERR599112 |
| TARA_064 | 5 | -29.5019 | 37.9889 | ERR598970, ERR599088, ERR599150 |
| TARA_064 | 65 | -29.5333 | 37.9117 | ERR598972, ERR599023, ERR599025 |
| TARA_064 | 1000 | -29.5046 | 37.9599 | ERR599021, ERR599164 |
| TARA_065 | 5 | -35.1728 | 26.2868 | ERR598979, ERR599146 |
| TARA_065 | 30 | -35.2421 | 26.3048 | ERR598990, ERR599018, ERR599110 |
| TARA_065 | 850 | -35.1889 | 26.2905 | ERR598960, ERR599034 |
| TARA_068 | 5 | -31.0266 | 4.665 | ERR599129, ERR599171, ERR599174 |
| TARA_068 | 50 | -31.027 | 4.6802 | ERR599017, ERR599056, ERR599103 |
| TARA_068 | 700 | -31.0198 | 4.6685 | ERR598947, ERR599131 |
| TARA_070 | 5 | -20.4091 | -3.1759 | ERR599135, ERR599165 |
| TARA_070 | 800 | -20.4075 | -3.1641 | ERR599044, ERR599149 |
| TARA_072 | 5 | -8.7789 | -17.9099 | ERR598984, ERR599105 |
| TARA_072 | 100 | -8.7296 | -17.9604 | ERR599133, ERR599137 |
| TARA_072 | 800 | -8.7986 | -17.9034 | ERR599005, ERR599048 |
| TARA_076 | 5 | -20.9354 | -35.1803 | ERR599010, ERR599126 |
| TARA_076 | 150 | -21.0292 | -35.3498 | ERR599040, ERR599148 |
| TARA_076 | 800 | -20.9315 | -35.1794 | ERR599000, ERR599154 |
| TARA_078 | 5 | -30.1367 | -43.2899 | ERR599006, ERR599022 |
| TARA_078 | 120 | -30.1484 | -43.2705 | ERR599046, ERR599101 |
| TARA_078 | 800 | -30.1471 | -43.2915 | ERR599124, ERR599159 |
| TARA_085 | 5 | -62.0385 | -49.529 | ERR599090, ERR599176 |
| TARA_085 | 90 | -62.2231 | -49.2139 | ERR599104, ERR599121 |
| TARA_085 | 790 | -61.9689 | -49.5017 | ERR599008, ERR599125 |
| TARA_098 | 5 | -25.8051 | -111.7202 | ERR599093, ERR599120 |
| TARA_098 | 188 | -25.826 | -111.7294 | ERR599042, ERR599079 |
| TARA_098 | 488 | -25.8076 | -111.6906 | ERR599071, ERR599085 |
| TARA_102 | 5 | -5.2529 | -85.1545 | ERR598943, ERR598978 |
| TARA_102 | 40 | -5.2669 | -85.2732 | ERR598962, ERR599007, ERR599168 |
| TARA_102 | 480 | -5.261 | -85.1678 | ERR599055, ERR599128, ERR599132 |
| TARA_109 | 5 | 1.9928 | -84.5766 | ERR598997, ERR599118 |
| TARA_109 | 30 | 2.0299 | -84.5546 | ERR598952, ERR599065, ERR599108 |
| TARA_109 | 380 | 2.0649 | -84.5546 | ERR598971, ERR599067 |
| TARA_110 | 5 | -2.0133 | -84.589 | ERR599039 |
| TARA_110 | 50 | -1.9002 | -84.6265 | ERR599014 |
| TARA_110 | 380 | -1.8902 | -84.6141 | ERR599020 |
| TARA_111 | 5 | -16.9601 | -100.6335 | ERR599077 |
| TARA_111 | 90 | -16.9587 | -100.6751 | ERR598961 |
| TARA_111 | 350 | -16.9486 | -100.6715 | ERR599086 |
| TARA_112 | 5 | -23.2811 | -129.3947 | ERR598954 |
| TARA_112 | 155 | -23.2189 | -129.4997 | ERR598957 |
| TARA_112 | 696 | -23.2232 | -129.5986 | ERR599072 |
| TARA_122 | 5 | -8.9971 | -139.1963 | ERR598992 |
| TARA_122 | 115 | -9.0063 | -139.1394 | ERR598948 |
| TARA_122 | 600 | -8.9729 | -139.2393 | ERR598999, ERR599033, ERR599083, ERR599096 |
| TARA_132 | 5 | 31.5213 | -158.9958 | ERR599142 |
| TARA_132 | 115 | 31.5168 | -159.046 | ERR598995 |
| TARA_132 | 550 | 31.528 | -159.0224 | ERR598980 |
| TARA_133 | 5 | 35.3671 | -127.7422 | ERR599052 |
| TARA_133 | 45 | 35.4002 | -127.7499 | ERR598942 |
| TARA_133 | 650 | 35.2698 | -127.7268 | ERR599115 |
| TARA_137 | 5 | 14.2035 | -116.6261 | ERR598989 |
| TARA_137 | 40 | 14.2075 | -116.6468 | ERR598987, ERR599070, ERR599099, ERR599147 |
| TARA_137 | 375 | 14.2025 | -116.6433 | ERR599015, ERR599076, ERR599127, ERR599152 |
| TARA_138 | 5 | 6.3332 | -102.9432 | ERR599030 |

| | | | | |
|---|---|---|---|---|
| TARA_138 | 60 | 6.3378 | -102.9538 | ERR599087 |
| TARA_138 | 450 | 6.3559 | -103.0598 | ERR599004, ERR599051, ERR599060, ERR599175 |
| TARA_142 | 5 | 25.5264 | -88.394 | ERR599136 |
| TARA_142 | 125 | 25.6168 | -88.4532 | ERR599100 |
| TARA_142 | 640 | 25.6236 | -88.45 | ERR598985 |
| TARA_145 | 5 | 39.2305 | -70.0377 | ERR598983 |
| TARA_145 | 590 | 39.2392 | -70.0343 | ERR599166 |
| TARA_146 | 5 | 34.6712 | -71.3093 | ERR598968 |
| TARA_146 | 640 | 34.6663 | -71.2907 | ERR599047 |
| TARA_149 | 5 | 34.1132 | -49.9181 | ERR598963 |
| TARA_149 | 740 | 34.0771 | -49.8233 | ERR598964 |
| TARA_152 | 5 | 43.6792 | -16.8344 | ERR599078 |
| TARA_152 | 25 | 43.7056 | -16.8794 | ERR599001 |
| TARA_152 | 800 | 43.7182 | -16.8714 | ERR598944 |
| ALOHA | 25 | 22.7500 | -158.0000 | SRR5002398 |
| ALOHA | 75 | 22.7500 | -158.0000 | SRR5002330 |
| ALOHA | 125 | 22.7500 | -158.0000 | SRR5002364 |
| ALOHA | 200 | 22.7500 | -158.0000 | SRR5002329 |
| ALOHA | 500 | 22.7500 | -158.0000 | SRR5002399 |
| ALOHA | 770 | 22.7500 | -158.0000 | SRR5002397 |
| ALOHA | 1000 | 22.7500 | -158.0000 | SRR5002331 |
| Izu-Bonin Trench | 0 | 29.1500 | 142.8012 | DRR092705 |
| Izu-Bonin Trench | 306 | 29.1500 | 142.8012 | DRR092706 |
| Izu-Bonin Trench | 505 | 29.1500 | 142.8012 | DRR092707 |
| Izu-Bonin Trench | 754 | 29.1500 | 142.8012 | DRR092708 |
| Izu-Bonin Trench | 1206 | 29.1500 | 142.8012 | DRR092709 |
| Izu-Bonin Trench | 2015 | 29.1500 | 142.8012 | DRR092710 |
| Izu-Bonin Trench | 3507 | 29.1500 | 142.8012 | DRR092711 |
| Izu-Bonin Trench | 5010 | 29.1500 | 142.8012 | DRR092712 |
| Izu-Bonin Trench | 7512 | 29.1500 | 142.8012 | DRR092713 |
| Izu-Bonin Trench | 9697 | 29.1500 | 142.8012 | DRR092714 |
| Japan Trench | 0 | 36.1050 | 142.7261 | DRR092715 |
| Japan Trench | 200 | 36.0640 | 142.7279 | DRR092716 |
| Japan Trench | 550 | 36.0640 | 142.7279 | DRR092717 |
| Japan Trench | 1000 | 36.0640 | 142.7279 | DRR092718 |
| Japan Trench | 3500 | 36.0640 | 142.7279 | DRR092719 |
| Japan Trench | 7000 | 36.1050 | 142.7261 | DRR092720 |
| Mariana | 0 | 11.3673 | 142.4181 | DRR092721 |
| Mariana | 203 | 11.6398 | 142.4294 | DRR092722 |
| Mariana | 502 | 11.6398 | 142.4294 | DRR092723 |
| Mariana | 2000 | 11.6398 | 142.4294 | DRR092724 |
| Mariana | 3971 | 11.3699 | 142.4301 | DRR092725 |
| Mariana | 7900 | 11.3699 | 142.4301 | DRR092726 |
| Mariana | 10899 | 11.3676 | 142.4241 | DRR092727 |
| Kuril Trench | 0 | 41.8736 | 146.3193 | DRR092728 |
| Kuril Trench | 200 | 41.8736 | 146.3193 | DRR092729 |
| Kuril Trench | 1000 | 41.8736 | 146.3193 | DRR092730 |
| Kuril Trench | 2400 | 41.8736 | 146.3193 | DRR092731 |
| Kuril Trench | 4500 | 41.8736 | 146.3193 | DRR092732 |
| Kuril Trench | 6500 | 41.8736 | 146.3193 | DRR092733 |

**Supplementary Table 2 | Cellular fatty acid composition of strain JH545 and representative species of the phylum *Chloroflexota*. Strains**: 1, “*Candidatus* Lucifugimonas marina” JH545 (this study); 2, *Dehalococcoides mccartyi* 195$^{T}$ [1]; 3, *Anaerolinea thermophile* UNI-1$^{T}$ [2]; 4, *Anaerolinea thermolimosa* IMO-1$^{T}$ [3]; 5, *Thermosporothrix hazakensis* SK20-1$^{T}$ [4]; 6, *Tepidiforma bonchosmolovskayae* 3753O$^{T}$ [5]. −, not detected.

| Fatty acids | 1 | 2 | 3 | 4 | 5 | 6 |
|---|---|---|---|---|---|---|
| **Saturated** | | | | | | |
| $C_{12:0}$ | 4.1 | − | 1.0 | 1.0 | − | − |
| $C_{14:0}$ | 4.3 | 15.7 | − | 5.0 | − | − |
| $C_{15:0}$ | − | − | 14.0 | 2.0 | − | − |
| $C_{16:0}$ | 10.1 | 22.7 | 35.0 | 16.0 | 10.0 | − |
| $C_{17:0}$ | − | − | 7.0 | 3.0 | 1.8 | − |
| $C_{18:0}$ | 2.6 | 16.6 | 12.0 | 3.0 | 7.3 | − |
| $C_{20:0}$ | − | − | − | − | − | 3.9 |
| **Unsaturated** | | | | | | |
| $C_{16:1}$ 2-OH | − | − | − | − | 9.4 | − |
| $C_{18:1}$ ω9 | − | − | − | − | 0.7 | − |
| **Branched chain** | | | | | | |
| anteiso-$C_{13:0}$ | − | − | − | 3.0 | − | − |
| iso-$C_{13:0}$ | − | − | − | 5 | − | − |
| brC 14:0 | − | − | 1.0 | − | − | − |
| brC 15:0 | − | 6.2 | − | − | − | − |
| anteiso-$C_{15:0}$ | − | − | − | 12.0 | − | − |
| iso-$C_{15:0}$ | − | − | − | 19.0 | 0.6 | − |
| 10-methyl $C_{16:0}$ | − | 25.8 | − | − | 1.3 | − |
| iso-$C_{16:0}$ | − | − | − | 2.0 | 1.1 | − |
| brC 17:0 | − | − | 6.0 | − | − | − |
| anteiso-$C_{17:0}$ | − | − | 6.0 | 21.0 | 10.3 | − |
| iso-$C_{17:0}$ | 4.3 | − | − | 5.0 | 52.8 | − |
| 10-methyl $C_{18:0}$ | 20.2 | − | − | − | − | − |
| brC 19:0 | − | − | 6.0 | − | − | 90.4 |
| brC 21:0 | − | − | − | − | − | 4.4 |
| **Summed feature 9*** | 54.4 | − | − | − | − | − |

*Summed features are groups of two or three fatty acids that are treated together for the purpose of evaluation in the MIDI system and include both peaks with discrete ECLs as well as those where the ECLs are not reported separately. Summed feature 9 comprises $C_{17:1}$ ω9c and/or 10-methyl $C_{16:0}$.

### Supplementary Text

#### Genome-based phylogeny of the SAR202 clade

Genome-based phylogeny showed that the entire SAR202 clade corresponds to a monophyletic superorder in the Genome Taxonomy Database (GTDB), with the SAR202 strains of this study belonging to the SAR202 group I, which corresponds to an order in the GTDB. To find the phylogenomic position of the four SAR202 strains in the context of both the previously proposed SAR202 groups (I to VII) and the GTDB[6-8], we classified SAR202 genomes (the four genomes of this study and other SAR202 MAG/SAGs reported previously) using GTDB-Tk, and then constructed a phylogenomic tree that encompasses GTDB taxa into which the SAR202 genomes were classified. The resulting tree (Fig. 1b) confirmed the phylogenetic position of the four strains analyzed by 16S rRNA gene phylogeny: the four strains belonged to the SAR202 group I, which corresponds to o__UBA1151 in the GTDB taxonomy (more specifically, group Ia corresponding to f__Bin127 in the GTDB R202). The whole SAR202 clade was revealed to be a monophyletic superorder (~10 orders) of the class *Dehalococcoidia* in the GTDB taxonomy, with most SAR202 groups corresponding to one of the orders, except for group V containing two orders. The order named "o__SAR202" in the GTDB is just one of multiple orders comprising the whole SAR202 clade. The orders SAR202-VII-2, GCA-2717565, and UBA1127 do not correspond to the groups of the SAR202 clade, but they are regarded to belong to the SAR202 clade, because these orders were nested within the orders corresponding to the SAR202 groups. The description of the SAR202 clade as a superorder is also consistent with a previous analysis, which showed a within-clade 16S rRNA gene sequence similarity as low as 78.7%[6], which is slightly higher than the threshold for class as proposed by Yarza *et al.*[9] This enormous phylogenetic diversity of the SAR202 clade may underlie metabolic and ecological divergence among the SAR202 groups reported previously[7].

**Proposal of ranks of the new taxa**

**Description of "*Candidatus* Lucifugimonas" gen. nov.**

*Lucifugimonas* (Lu.ci.fu.gi.mo'nas. L. fem. n. *lux*, lucis, light; L. fem. n. *fuga*, flight; L. fem. n. *monas*, a unit, monad; N.L. fem. n. *Lucifugimonas*, a monad that prefers dark habitats).

Aerobic, oligotrophic, and chemoheterotrophic. Gram-negative with a monoderm envelope. Cells are non-motile and dimorphic with short rods of ~0.8 × 0.4 µm and cocci of ~0.5 µm diameter. Do not form colonies on solid agar plates. Grows very slowly and reaches stationary phase at ~50 days of growth in artificial seawater medium. Light inhibits cellular growth. Major cellular fatty acids are summed feature 9 ($C_{17:1}$ ω9c and/or 10-methyl $C_{16:0}$), 10-methyl $C_{18:0}$, and $C_{16:0}$, different from other representatives of *Chloroflexota* (Supplementary Table 2). The genus "*Candidatus* Lucifugimonas" is assigned to SAR202 group Ia within the class *Dehalococcoidia* based on 16S rRNA gene phylogeny and whole genome phylogenomics. The type species of the genus is "*Candidatus* Lucifugimonas marina".

**Description of "*Candidatus* Lucifugimonas marina" sp. nov.**

*Lucifugimonas marina* (ma.ri'na. L. fem. adj. *marina*, marine, of the sea).

In addition to the properties given in the genus description, the species is described as follows. Growth occurs at temperatures between 10 °C and 25 °C, but not at 4 °C or below, nor at 30 °C or above. Optimum growth temperature is 15−20 °C. Grows only in seawater-based liquid medium or artificial seawater medium. Cellular growth is enhanced by fucose, fuconate, fucono-1,4-lactone, rhamnose, rhamnono-1,4-lactone, rhamnonate, and ascorbate. The type strain, JH545, was isolated from epipelagic seawater off the coast of Garorim Bay at Tea-An, South Korea. The length of the complete whole genome sequence of the type strain is 3.08 Mbp with 51.8% of the DNA G+C content. GenBank accession number of the type strain is CP046146. Besides the type strain, whole genome sequences of strains JH639, JH702, and JH1073 belonging to this species are also available under GenBank accession numbers WMBD00000000, WMBE00000000, and CP046147, respectively.

**Description of "*Candidatus* Lucifugimonadaceae" fam. nov.**

*Candidatus* Lucifugimonadaceae (Lu.ci.fu.gi.mo.na.da.ce'ae. N.L. fem. n. *Lucifugimonas*, a bacterial genus; -*aceae*, ending to denote a family; N.L. fem. pl. n. *Lucifugimonadaceae*, the *Lucifugimonas* family).

The description is the same as with the genus "*Candidatus* Lucifugimonas". The type genus is "*Candidatus* Lucifugimonas". Equivalent to GTDB f__UBA1328 (R207).

**Description of "*Candidatus* Lucifugimonadales" order. nov.**

*Candidatus* Lucifugimonadales (Lu.ci.fu.gi.mo.na.da'les. N.L. fem. n. *Lucifugimonas*, a bacterial genus; -*ales*, ending to denote a family; N.L. fem. pl. n. *Lucifugimonadales*, the *Lucifugimonas* order)

The description is the same as with the genus "*Candidatus* Lucifugimonas". The type genus is "*Candidatus* Lucifugimonas". Equivalent to GTDB o__UBA1151 (R207).

**Genomic inference of the metabolic potentials**

The JH545 genome exhibits features of a heterotrophic lifestyle (Fig. 3), in accordance with previous studies on SAR202 MAG/SAGs. The genome has genes for central carbon and energy metabolism of typical aerobic organoheterotrophs, including the Embden-Meyerhof-Parnas glycolytic pathway, the tricarboxylic acid (TCA) cycle, oxidative and non-oxidative pentose phosphate pathway (PPP), and electron transport chain (except for cytochrome *c* reductase). Gluconeogenesis is incomplete due to the lack of fructose-1,6-bisphosphatase. Several features found in central metabolic pathways are as follows. In accordance with a previous study on SAR202 MAGs[10], decarboxylation of pyruvate to acetyl-CoA is mediated by both pyruvate dehydrogenase complex and pyruvate:ferredoxin oxidoreductase. In contrast to pyruvate, 2-oxoglutarate, a key intermediate of TCA cycle, is decarboxylated by only 2-oxoacid:ferredoxin oxidoreductases. Alpha-ketoglutarate (2-oxoglutarate) dehydrogenase complex was not predicted in the genomes. The first step of oxidative PPP is mediated by glucose-6-phosphate dehydrogenase ($NAD^+$) (EC 1.1.1.388), instead of more generally used glucose-6-phosphate dehydrogenase ($NADP^+$) (EC1.1.1.49). The enzymes assigned to EC 1.1.1.388 belong to a novel type of glucose-6-phosphate dehydrogenase discovered in a recent study that demonstrated oxidative PPP for the first time in the domain Archaea[11]. This novel enzyme group, assigned to K19243 in the KEGG database, is not related to the enzymes belonging to EC 1.1.1.49 (K00036). A search for K19243 proteins in representative genomes for species clusters of the GTDB (R95), conducted using AnnoTree, revealed that 3.8% (1,135 of 30,238) and 18.2% (304 of 1,672) of bacterial and archaeal genomes possessed this novel enzyme, respectively. While rare in bacteria, this novel enzyme is found in all genomes of o__UBA1151, suggesting the importance of this enzyme in carbon metabolism of SAR202 subgroup I.

Like many marine heterotrophs[12], the SAR202 genomes were predicted to be deficient in the biosynthetic pathways of several vitamins, including thiamine ($B_1$), biotin ($B_7$), and cobalamin ($B_{12}$). The lack of thiamine phosphate synthase, in addition to the biosynthetic pathways of the precursors, suggested a requirement for intact $B_1$. Only a salvage pathway was found for cobalamin. Folate ($B_9$) biosynthesis is also questionable due to the absence of several steps including those for biosynthesis of 4-aminobenzoate, a key precursor of folate. Also, it was predicted that the biosynthetic pathways for serine and cysteine could be incomplete due to the lack of *serB* (phosphoserine aminotransferase) or *cysE* (serine O-acetyltransferase), which would require wet experiments for verification.

In addition to genes functioning in aerobic heterotrophic metabolism, some genes involved in lithotrophy and anaerobic respiration were also found in the JH545 genome. Strain JH545 is predicted to use sulfide as electron donor. Sulfide:quinone oxidoreductase (EC 1.8.5.4), which directly transfers electrons from sulfide to the quinone pool, was predicted to be present in the genome, suggesting the possibility of utilizing sulfide as an energy source[13]. In contrast to previous studies[10,14], genes necessary for energy conservation by oxidation of sulfite or methanesulfonate were not found. Although lithoautotrophy is unlikely due to the absence of any carbon fixation pathways, the energy from sulfide would contribute to survival in oligotrophic habitats. In terms of electron acceptors, nitrate reductase (NapAB) and $N_2O$ reductase (NosZ) were predicted. Although no energy conservation is possible since both enzymes are periplasmic, utilization of these nitrogen compounds as electron acceptors may provide an advantage in anaerobic conditions that may be encountered in some habitats. The biosynthetic pathway for molybdenum cofactor (Moco), an essential coenzyme of nitrate reductase, and the ABC transporter for molybdate were also predicted to be encoded in the genome.

### Supplementary Method

#### Analysis of fatty acid composition

The analysis of cellular fatty acid methyl esters (FAMEs) was performed using the standard protocol provided by the MIDI/Hewlett-Packard Microbial Identification System[15]. To extract FAMEs, cells of strain JH545 were harvested by centrifugation at 13,000 g for 1 hr at the end of the exponential growth phase in a 400 mL volume of liquid culture (ASW5x media). The extracted FAMEs were saponified and methylated before being analyzed using a gas chromatograph (Agilent 7890 GC) with TSBA6 database from the Sherlock Microbial Identification System (MIDI) version 6.1.
